# Binding of SARS-CoV-2 nucleocapsid protein to uninfected epithelial cells induces antibody-mediated complement deposition

**DOI:** 10.1101/2024.03.17.585388

**Authors:** Jamal Fahoum, Maria Billan, Julia K Varga, Dan Padawer, Maya Elgrably-Weiss, Pallabi Basu, Miri Stolovich-Rain, Leah Baraz, Einav Cohen-Kfir, Sujata Kumari, Esther Oiknine-Dijan, Manoj Kumar, Orly Zelig, Guy Mayer, Michail N Isupov, Dana G Wolf, Shoshy Altuvia, Reuven Wiener, Ora Schueler-Furman, Alexander Rouvinski

## Abstract

SARS-CoV-2 infection triggers strong antibody response toward Nucleocapsid-Protein (NP), suggesting extracellular presence beyond its intra-virion RNA binding. Interestingly, NP was found to decorate infected and proximal uninfected cell-surfaces. Here, we propose a new mechanism through which extracellular NP on uninfected cells contributes to COVID-19 pathogenicity. We show that NP binds to cell-surface sulfated linear-glycosaminoglycans by spatial rearrangement of its RNA-binding sites facilitated by the flexible, positively charged, linker. Coating of uninfected lung-derived cells with purified NP attracted anti-NP-IgG from lung fluids and sera collected from COVID-19 patients. The magnitude of this immune recognition was significantly elevated in moderate compared to mild COVID-19 cases. Importantly, binding of anti-NP-IgG present in sera generated clusters that triggered C3b deposition by the classical complement pathway. Heparin analog enoxaparin outcompeted NP-binding, rescuing cells from anti-NP IgG-mediated complement deposition. Our findings unveil how extracellular NP may exacerbate COVID-19 tissue damage, and suggest leads for preventative therapy.

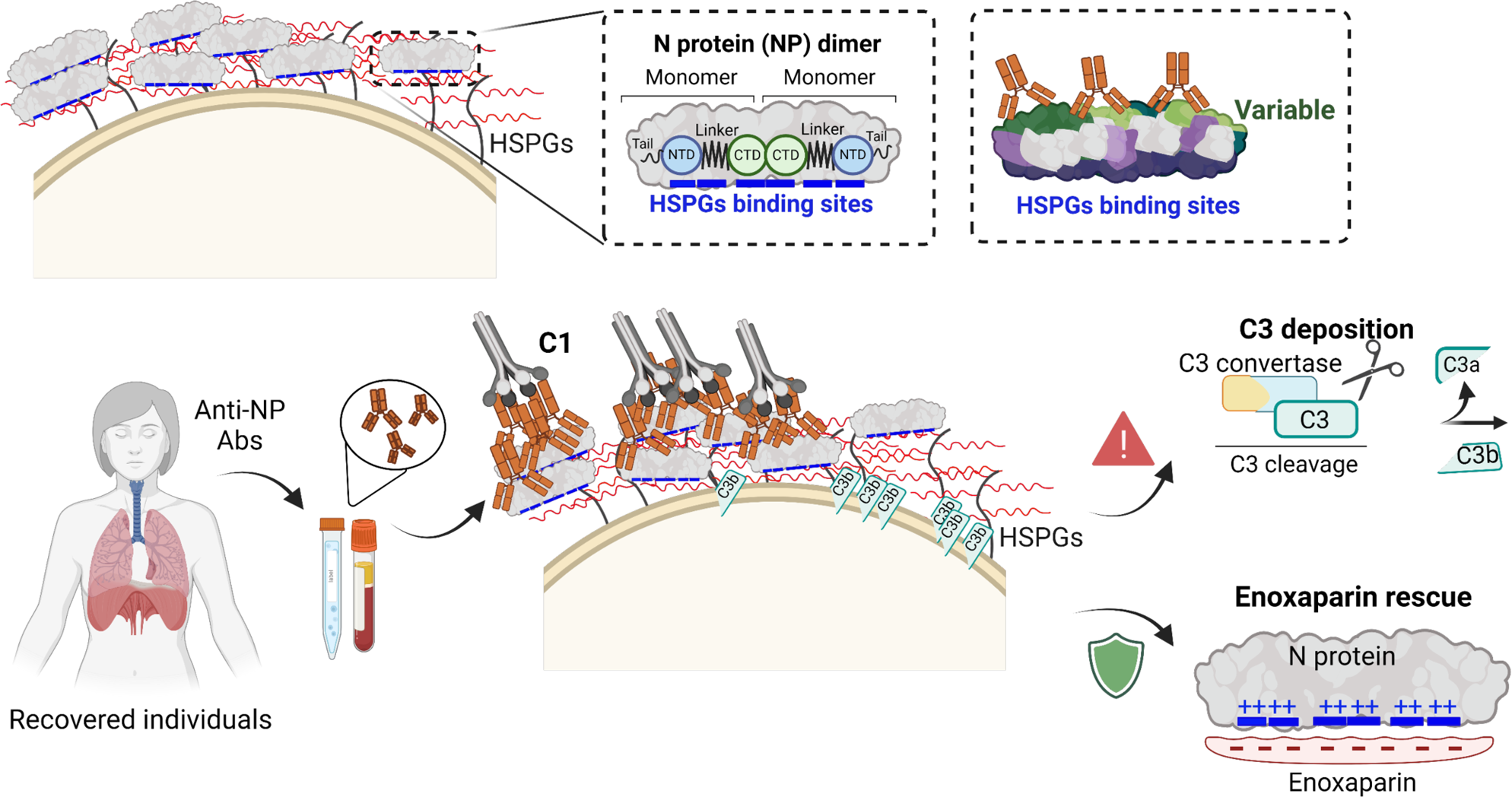

**Highlights:**

- IgG from patients’ sera target NP-bound cells resulting in complement activation
- The flexible linker allows NP to both bind linear sulfated GAGs and wrap around RNA
- Heparin analogs prevent NP surface binding and alleviate complement activation
- Cell-ELISA anti-NP IgG levels differ between mild and moderate COVID-19

## Introduction

Since the outbreak of severe acute respiratory syndrome coronavirus 2 (SARS-CoV-2) in 2019 in Wuhan, China, hundreds of millions of people around the world have been infected. While the majority fully recover, a wide spectrum of clinical manifestations has been reported, including life threatening disease in viral and post viral phases and long term post COVID-19 syndromes (1–3). The post viral stages of the disease are often associated with a strong, immune system mediated, cytokine storm with a fraction of patients experiencing long-term post-recovery symptoms, known as long COVID. Those signs vary from mild to more severe, involving long–term tissue damage (3) such as lung fibrosis (2,4–6). Severe COVID-19 cases are more likely to develop long-term post COVID-19 symptoms (3). However, the risk factors and biomarkers associated with developing disease severity and post COVID-19 await to be fully deciphered.

Several recent reports suggest that while the prevalence of spike-specific antibodies is common in mild cases, high titers of nucleocapsid protein (NP) - specific antibodies, both IgM and IgG, correlate with disease severity (7,8), whereas upon vaccination NP antibodies were shown to be protective (9). A recent study has clearly shown physiological secretion of NP to the surface of infected cells (10,11). Surface exposure of NP might target infected cells for detection by the immune system. Importantly, NP can rapidly spread by association with cell surface heparan sulfate proteoglycans (HSPGs) from infected to nearby uninfected cells, including cells resilient to infection (10). Indeed, in addition to RNA binding, NP was shown to bind to a soluble heparin analog, enoxaparin, as recently demonstrated by Nuclear Magnetic Resonance (NMR) (12). Cellular HSPGs often exhibit co-receptor roles due to their ability to bind multiple host ligands, receptors and signaling molecules in the extracellular space, as well as heterologous ligands, such as viruses through the sulfated polymeric glycosaminoglycans (GAGs) attached to serine residues of the protein core. Due to their long-distance electrostatic potential and polymeric nature, HSPGs can attract ligands, stabilize them and assist their lateral distribution to neighboring cells (13).

The existence of NP on the cell surface of uninfected cells (10) raises a possibility of skewed immune response triggered by Fc of anti-NP IgGs, via activation of the classical complement pathway (14). Antibody mediated complement activity is based on targeting of pathogens by specific antibodies and induction of the cascade activation pathway of complement factors that recognize and bind clustered Fc regions of the antibodies (14,15). In a well-balanced scenario, confinement of the infection zone including sacrificing proximal cells (in particular those susceptible to infection) should isolate and restrict the infection spread. However, in exaggerated cases, such targeting could lead to uncontrolled activation of the immune system toward cells with surface bound NP and lead to severe tissue damage which could be the cause for severe COVID-19 and post COVID-19 pathologies. Moreover, such retargeting may partially subvert the immune response from virus infected cells toward uninfected neighboring cells.

The molecular features of NP that determine its binding properties to the cell surface are not known. It is not clear how NP could bind also to linear, extended GAG chains, such as those of HSPGs on the cell surface, and whether this binding mode impacts its interplay with the immune system (see e.g. (16)). Intriguingly, despite multiple studies on NP biochemical and structural characteristics, its modalities of RNA association are not yet fully explored. NP consists of two structured domains, the N-terminal domain (NTD), and the C-terminal domain (CTD), which are connected by a linker. The CTD serves as a dimerization domain of NP, which in turn has also been reported to form higher order oligomers via its linker region (17), as well as via its C-terminal helix (18). Many previous studies have demonstrated the importance of NTD in association with viral RNA and described the atomic details of these interactions (19). Several reports indicate possible roles of CTD in RNA binding as well, though atomic resolution structures of a CTD-RNA complex are yet lacking (19). Based on experimental crosslinks, a structural model of NP that includes two NTD monomers and one CTD dimer wrapping around RNA was generated (20). It remains to be determined to which extent do cell-surface determinants overlap with those involved in RNA packaging, given the significant similarity of the electrostatic potential of cell-surface sulfated proteoglycans and nucleic acids.

In this study, we elucidate the immuno-regulatory roles played by NP in SARS-CoV-2 pathogenicity, beyond RNA binding and packaging. Using a range of different experimental approaches, structural modeling, and a well-characterized pre-vaccination ‘era’ COVID-19 patient cohort, we decipher step-by-step the details of how NP binding to cells elicits decoration by anti-NP antibodies and subsequent complement activation. We determine the contribution of different parts of NP to binding to the cell surface, and model and experimentally confirm the structural basis of NP binding to HSPGs. We show that anti-NP IgG from human fluids recognize cell surface NP clusters and elicit complement cascade activation, as monitored by C3b deposition on uninfected NP-labeled cultured cells. Using samples and clinical data from our cohort, we characterize the association of anti-NP IgG with COVID-19 severity, and reveal that COVID-19 complications, such as lung radiological abnormalities, are associated with high levels of anti-NP IgG in serum capable to recognize epitopes presented by cell surface bound NP.

We propose a scenario, whereby the complement targets uninfected cells marked by an exogenously associated pathogen attribute, subverting the immune response from virus propagating cells to the uninfected neighboring cells. Furthermore, we show that enoxaparin, a low-weight heparin analog, prevents binding of NP to the cell surface and thereby rescues NP-coated cells from antibody-mediated complement deposition, opening a new avenue for the prevention of the NP-driven immunological complications of COVID-19.

## Results

### Antibodies against NP are found in sera and Broncho-Alveolar Lavage (BAL) fluid of COVID-19 recovered patients

While NP is mainly considered an intracellular or intra-virion protein, multiple studies have reported anti-NP antibodies in the serum of recovered patients (8,21). Accordingly, in our analysis of sera from a patient cohort, we observed the presence of anti-NP IgG in the serum of patients that recovered from SARS-CoV-2 infection in the pre-vaccination era of the pandemic, but not in the serum of a corresponding naïve pre-COVID-19 population (ELISA assay, **Figure 1A**). Moreover, we also detected IgG against NP in a cohort of 16 patients that had undergone bronchoscopy. Among these, COVID-19 recovered patients showed antibodies against NP not only in their serum (**Figure 1B**), but also in the BAL fluid (**Figure 1C**), pointing to the possibility of NP involvement in alveolar pathology of COVID-19 patients. Information about the different cohorts used in this study is summarized in **Supplementary Table S1**.

**Figure 1.**
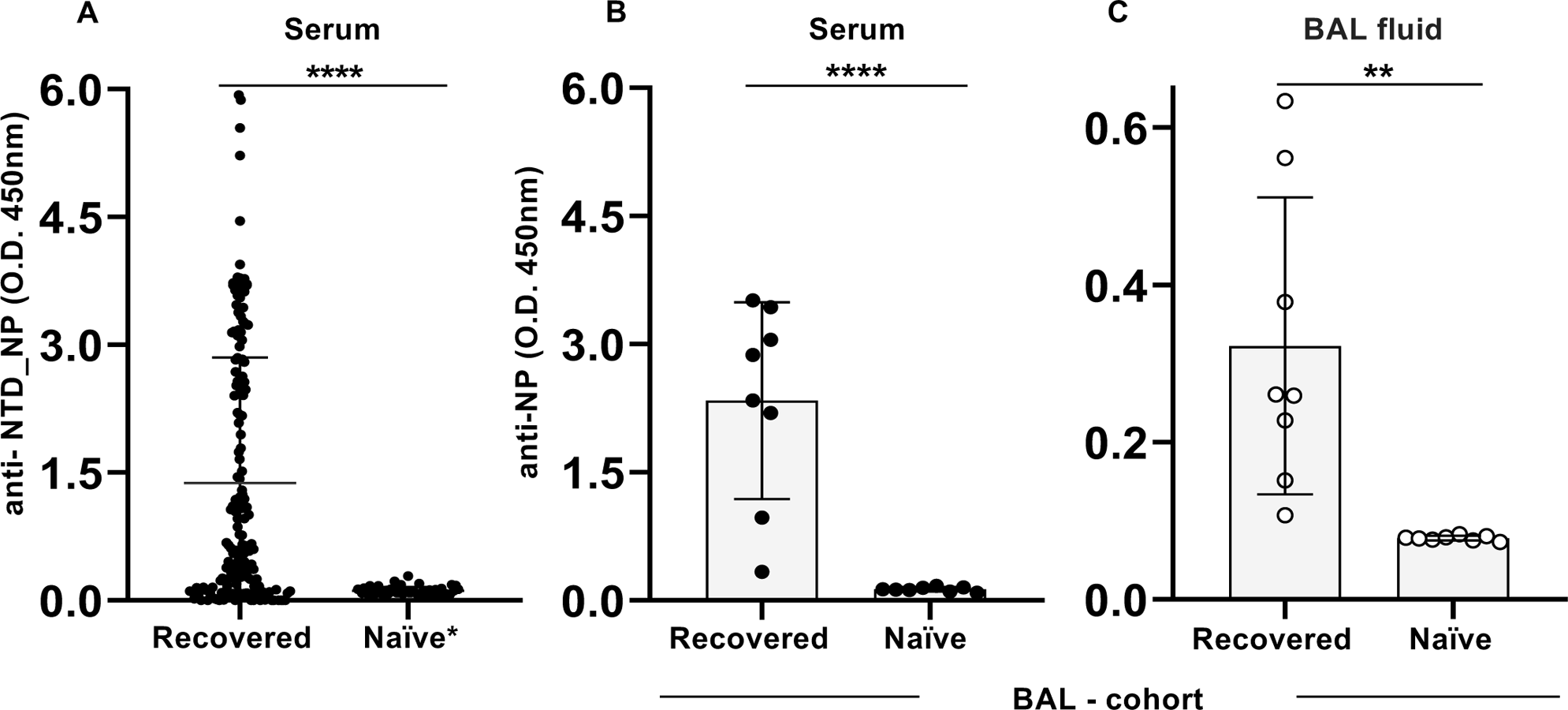
Antibodies against NP are found in sera and bronchoalveolar lavage (BAL) of COVID-19 recovered individuals. **A:** ELISA detection of anti-NTD_NP antibodies in sera collected from recovered COVID-19 cases (n=170) and naïve (n=51) individuals. Naïve* indicates samples collected in the pre-COVID-19 era. **B, C:** ELISA for detection of anti-NP antibodies in a cohort of individuals that underwent BAL, both in their serum (**B**) and BAL fluids (**C**) samples, including COVID-19 recovered (n=8) and naïve (n=8) individuals. Student’s two-tailed unpaired t-test was used to compare recovered *vs* naïve samples.’\**\*’* indicates *P value<*0.01, and ‘****’ indicates *P value<*0.0001.

### NP binds to the cell surface via sulfated proteoglycans

What could be the biological function of extracellular NP, in addition to the recently suggested cytokine scavenging (11) or the RAGE/MAPK (mitogen-activated protein kinase)/NF-ĸB pathway activation (16)? To explore this question, we expressed, purified and labeled NP with fluorescein isothiocyanate (FITC) using sortase transpeptidation (**Figure 2A,B**; see **Methods**) (22–24). Full-length, labeled NP binds to the cell surface of HeLa cells, as well as of A549 lung carcinoma epithelial cells, in a concentration dependent manner (as examined by flow cytometry assays, FACS, **Figure 2C**, and fluorescence microscopy, **Figure 2D**), in agreement with a recent report (10). While both HeLa and A549 cell lines show NP binding, the latter (originating from lung epithelium) displays significantly stronger binding capacity, suggesting either an increased affinity or a higher number of cell surface binding sites (**Figure 2C**). NP binding to the surface of these uninfected cells was also revealed by a western blot of cell extracts using his-tagged NP, detected by anti-his antibodies (**Figure 2E**). Furthermore, unlabeled NP was able to outcompete binding of both FITC-labeled and his-tagged NP, demonstrating specific binding (**Figures 2D,E**). A recent report suggested HPSGs as the most probable cellular receptor, supported by the absence of NP binding to CHO-pgsA-745 cells that lack HSPGs (10). In line with this finding, NP binding to HeLa cells is outcompeted by addition of pentosan polysulfate (PPS) that mimics soluble heparin (revealed by both fluorescence microscopy and western blots, **Figures 2F,G**).

**Figure 2:**
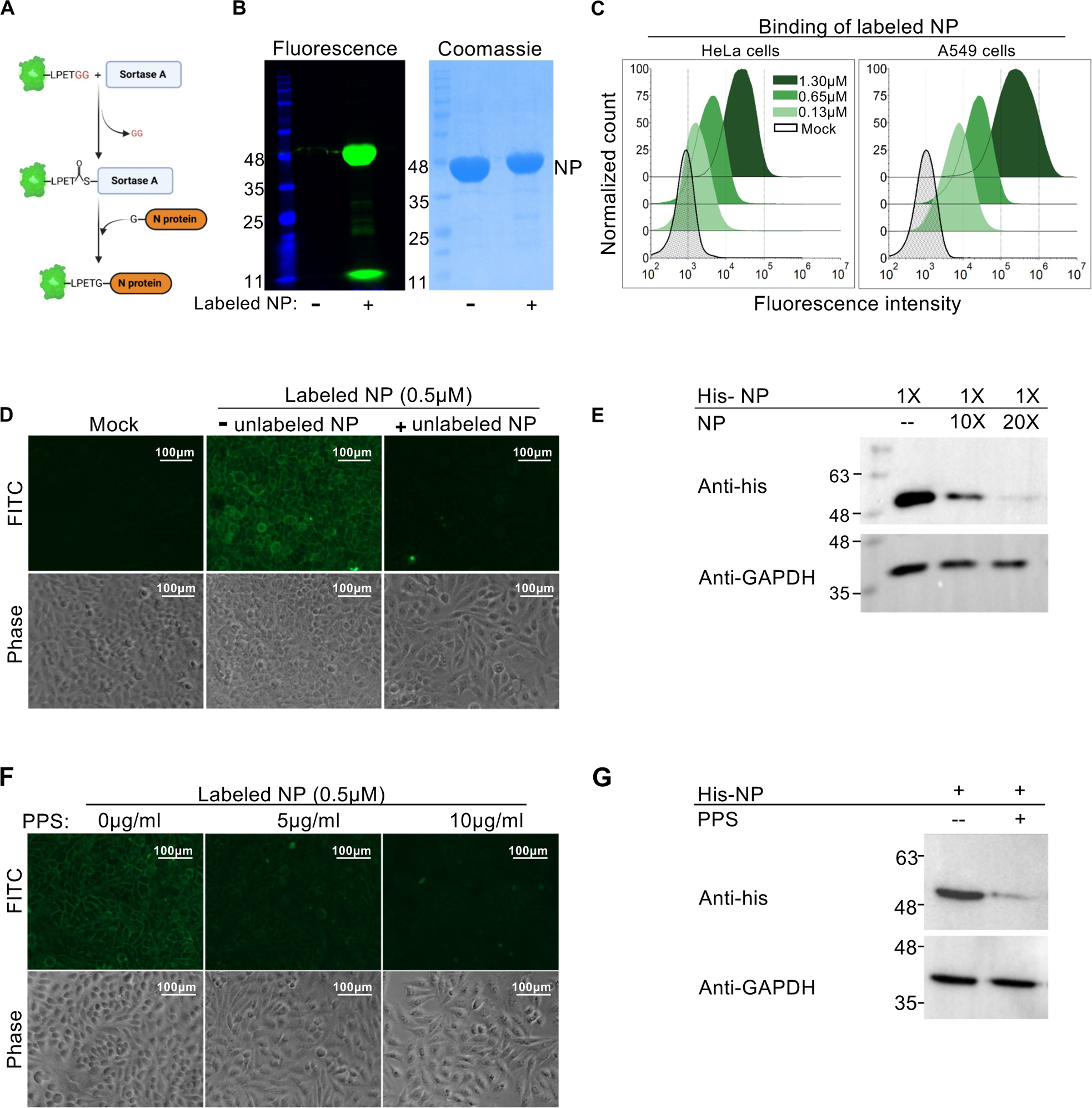
NP binds to the plasma membrane of uninfected cells. **A:** Schematic representation of N-terminal sortase mediated fluorescence labeling of NP by transpeptidation reaction. **B:** SDS-PAGE analysis of NP and its FITC-labeled derivative (left and right lane, respectively), imaged in fluorescence mode (left panel), (ex488nm/em520nm) and in transmitted light mode after Coomassie blue stain (right panel). Molecular weight (MW) values are indicated on the left side in kDa. **C:** Flow cytometry analysis of live HeLa and A549 cells incubated with FITC-labeled NP shows dose-dependent binding. The gating strategy is described in **Supplementary Figure S1A**. **D:** FITC-labeled NP was added to live HeLa cells (at 0.5μM concentration, middle panel; see **Supplementary Figure S1B** for additional concentrations). Binding is outcompeted after re-incubation with unlabeled NP (right panel). **E**: Reversible NP binding to the cell surface of HeLa cells revealed by a displacement assay. His-tagged NP was first bound to the surface of HeLa cells. Untagged NP in 10- and 20-fold excess amounts was added to cells for displacement assay. Western blot (WB) analysis of cell extracts with anti-his antibody is presented. **F-G:** Pentosan Polysulfate (PPS) outcompetes NP binding to cells. **F:** FITC-labeled NP (0.5μM) was incubated with live HeLa cells. Cells were washed and subsequently incubated with PPS at the indicated concentrations, and visualized by fluorescence microscopy. **G:** His-tagged NP was incubated with HeLa cells in either presence or absence of PPS, as indicated, and anti-his antibodies were used to detect the reduction of bound NP in cell extracts by WB. Scale bar represents 100μm. All experiments were performed at 4°C, to prevent endocytosis.

### The structural determinants of NP binding to cell surface, heparin and RNA

NP contains two defined domains: the N-terminal and the C-terminal domains (NTD and CTD, respectively; **Figure 3A**). NTD is well characterized for its binding to RNA (25), and RNA binding has also been demonstrated for CTD (e.g. (26)). These two globular domains are flanked by flexible tails and are connected by a linker. To characterize the HSPG binding sites, we modeled the interaction of heparin with the two structured NP domains, using a global docking server developed for the identification of heparin binding sites on proteins (27), supplied with solved crystal structures for NTD (Protein Data Bank, PDB, ID 6VYO (28,29)) and CTD (8R6E, a structure solved by us, see **Supplementary Figure S2** and **Table S2**). When heparin is docked to the individual domains, it occupies the RNA binding site on the NTD and the positively charged patch on the CTD that is most probably also used to bind RNA (20,26,30) (**Figure 3B**, all structural models are provided as **Supplementary Data**). These sites are also the most conserved in NP (**Figure 3A**). While a high resolution, confident structure of full-length NP has not been reported yet, a recently published cryoEM structure of full NP suggests an extended conformation (PDB ID 8FD5 (31)), in which the positive patches of the CTD dimer and the two NTD domains are all located on the same side and thus appear to be well positioned to bind the cell surface HSPGs (**Figure 3C**, left panel). Moreover, docking of heparin to this structure identified yet another potential binding site that is contributed by the positively charged interdomain linker (**Supplementary Figure S2C** and **Figure 3C**, left panel). While quality and reliability of this structure have been questioned by many researchers (see comment in PubPeer (32)), the suggested overall spatial organization of the protein may provide an explanation of how NP could strongly bind to linear sulfated GAG chains of cell surface HSPGs. Of note, this structure suggests a very different relative orientation of the NTD and CTD domains in comparison to a previously proposed structural model of NP bound to RNA, in which NP tightly wraps around the RNA to protect it (20) (**Figure 3C**, right panel). This highlights the importance of the interdomain flexible linker (see **Discussion**).

**Figure 3:**
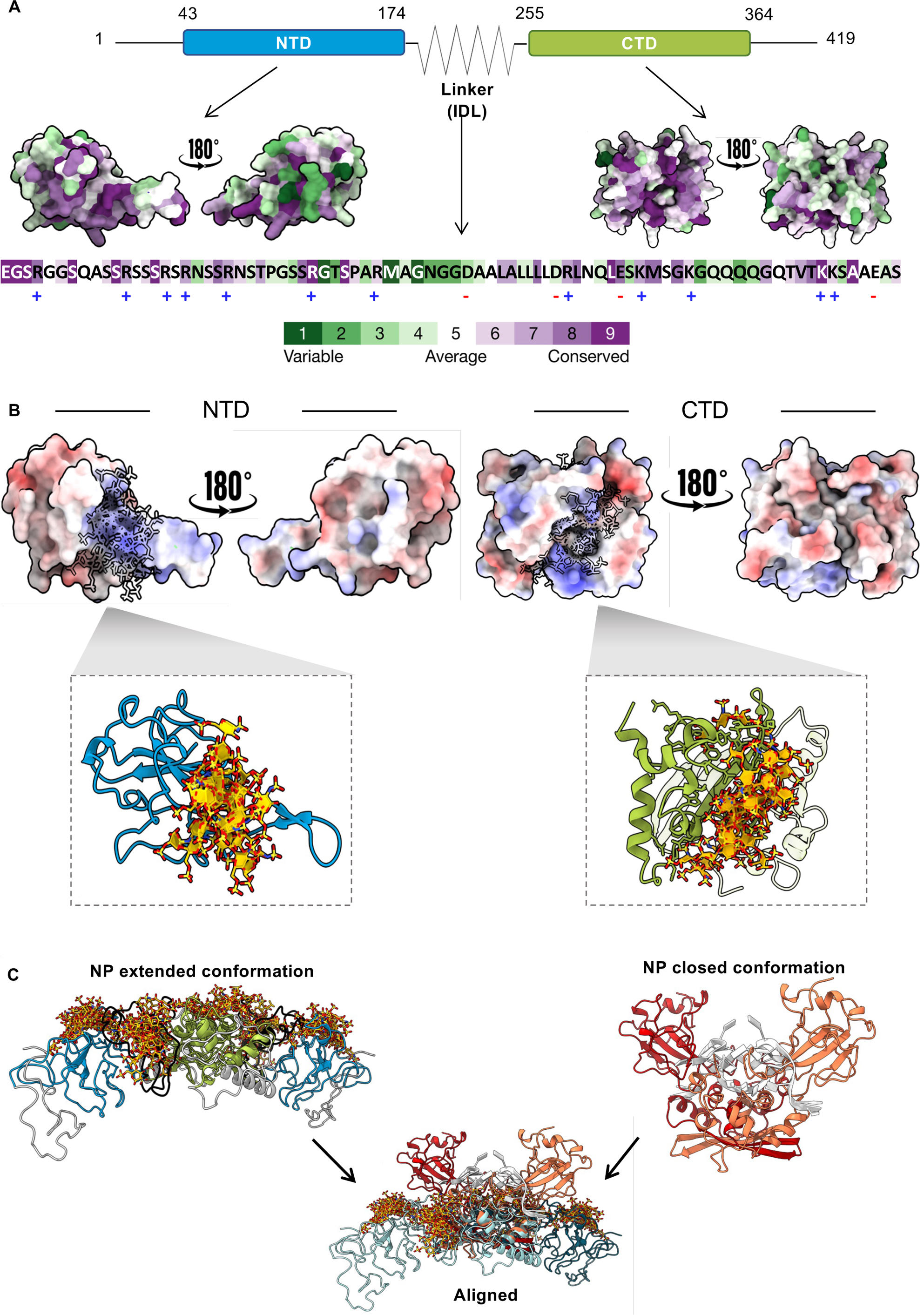
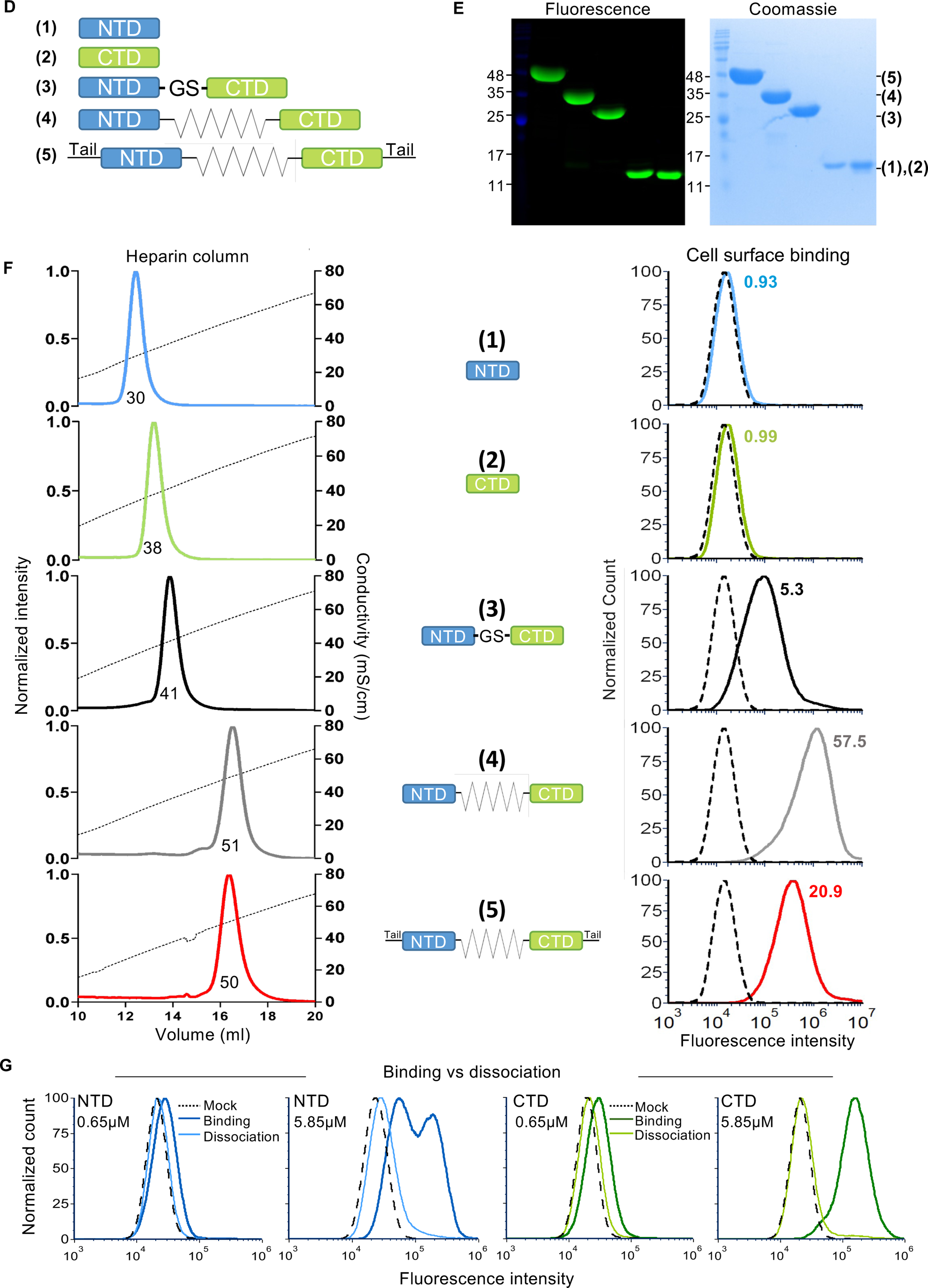
Contribution of different parts of NP to binding to heparin and to the cell surface. **A:** Architecture of NP: Two structured domains, the N-terminal Domain (NTD), and the C-terminal domain (CTD) are connected by a flexible interdomain linker (IDL), and flanked by flexible N-terminal and C-terminal tails. The linker sequence is shown colored according to conservation (determined using ConSurf (69)), and charged residues are indicated below the amino acid sequence to emphasize the overall positive charge. Structural models of NTD and CTD are also colored according to sequence conservation. **B:** Structural models of NTD (left panel) and CTD (dimer, right panel) bound to heparin (an ensemble of 8 models is shown). **Top:** The domains are colored according to their electrostatic potential, highlighting the positive patch at the binding site. Two different orientations are depicted, as indicated. Note the high degree of sequence conservation of the binding site, evident by comparison to the ConSurf representation of the NTD and CTD domains in panel A. **Bottom:** Cartoon representation of models of NTD (blue) and CTD (green) bound to heparin (yellow). **C:** Structural model of the extended, full-length dimer NP conformation bound to heparins. NP is shown in cartoon representations, colored according to regions: NTD (blue), IDL (black) and the CTD dimer (green). **Left:** side view, highlighting the extended heparin binding surface on the top. **Right:** RNA bound conformation of NP (see Figure 4 in reference (20)). **Middle:** Alignment between the structure in the extended, heparin bound conformation (top, left) and the closed, RNA bound conformation (top, right). **D:** Schematic representation of NP and the five constructs studied in the current work: (1) NTD, (2) CTD, (3) NTD-GS-CTD (short artificial interdomain linker, no tails), (4) NTD-IDL-CTD (missing the terminal tails), and (5) wt full length NP. **E:** SDS-PAGE of the five FITC labeled constructs, imaged in fluorescence mode (left) and after Coomassie blue staining (right). MW marker values are indicated in kDa. **F: Left panels:** Elution chromatograms of the five constructs from a heparin column (1ml) using NaCl gradient. The values under the curves represent conductivity (mS/cm) at the elution peak of the corresponding construct. Linear dashed diagonal lines represent conductivity change along the run due to NaCl gradient rise (right Y axis). The normalized absorbance (at 280nm) indicating the protein distribution is displayed as a solid line. **Right panels:** Flow cytometry analysis of live A549 cells incubated with the five FITC-labeled NP constructs (0.65μM). The dashed peak represents background control (mock, no NP protein, cells only). Median fluorescence intensity values for each peak are shown (x10^4^). **G:** Flow cytometry analysis of live A549 cells incubated with FITC-labeled NP domains NTD (left panels, blue curves) and CTD (right panels, green curves) at two experimental setups: binding (dark lines) *versus* dissociation (bright lines) for comparison. Each experiment was performed at two protein concentrations (0.65μM) and (5.85μM), as indicated. **Binding:** cells were only briefly washed once prior to flow cytometry analysis, while in **Dissociation:** cells were washed three times and were allowed to reach equilibrium for 10 min prior flow cytometry analysis. Experiment was performed at 4°C, to prevent endocytosis.

In order to experimentally verify the role of the different NP regions in cell surface and RNA binding, we generated and labeled five different NP constructs (**Figure 3D,E**): the full protein (Full), NTD, CTD, a construct that lacks the flexible tails (NTD_IDL_CTD, IDL=Inter-Domain Linker), and a construct that in addition lacks the native linker (NTD_GS_CTD; the IDL is replaced by a 14 residue linker, containing a thrombin cleavage site flanked by GSGS sequences: GSGS<u>LVPRGS</u>GSGS). We used this set of constructs to quantitatively compare NP binding to cell surface HSPGs *versus* RNA. First, we measured the ionic strength required for displacement of NP constructs from a heparin column, to rank their binding strength to heparin (**Figure 3F**, left panels). We noticed that CTD binds stronger than NTD to the heparin column, while all constructs containing both domains bind significantly stronger than the isolated domains. Moreover, while removing the tails of full NP did not significantly affect binding, replacing the IDL (in NTD_GS_CTD) reduced heparin binding, highlighting the importance/contribution of the native linker for binding to heparin, probably due to its significant positive charge (**Figure 3A**). Next, we measured the binding modalities of the NP constructs to the cell surface (**Figure 3F**, right panels). Using standard FACS experimental conditions, which included post-binding washes, we observed binding to the cell surface only in NP constructs containing both NTD and CTD. This discrepancy with the heparin binding results, which showed binding of NTD or CTD alone, prompted us to investigate whether the absence of binding in the FACS assay by individual NTD and CTD domains was due to the rapid dissociation of the single domains from the cell surface. For this purpose, we conducted an additional set of FACS experiments without extensive washing. Indeed, in the absence of washes, both NTD and CTD clearly demonstrated binding to the cell (**Figure 3G**). Our data collectively suggest that both NTD and CTD are capable of binding to the cell surface individually, but that stable binding is achieved only in the presence of both domains linked together, most likely due to increased avidity. Moreover, the natural interdomain linker with its conserved positively charged residues (**Figure 3A**) increases binding significantly more than the shortened synthetic neutral 14 residue linker (see **Discussion**).

The finding that CTD binds more tightly than NTD to heparin is surprising, as it is the NTD that has been reported to be a major RNA binding determinant (33). To explore this further, we evaluated the ability of the different constructs to bind RNA. We first applied on-column urea treatment during IMAC affinity purification to remove bacterial RNA/DNA contaminants from NP (see **Methods**). This extra purification step was necessary, since NP co-purifies from bacterial expression with bacterial RNA and forms oligomers. The nucleic acid removal step resulted in a mobility shift on the SEC column, indicating oligomer disassembly (**Figure 4A**). Of note, all purified proteins in the current study underwent this urea wash procedure. Reassuringly, while before this treatment NP was unable to bind a 242 nucleotides long RNA in an EMSA assay, urea treated, nucleic acid free NP readily bound RNA and formed high molecular weight oligomers (**Figure 4B, lanes 2-4** *versus* **5-7**). Following this functional validation, we proceeded to test the partial constructs. NTD-IDL-CTD that lacks the flexible tails bound in a similar way as the full-length NP, indicating that the flexible tails do not significantly contribute to RNA binding (**Figure 4B, lanes 11-13** *versus* **5-7**). However, removing the positively charged interdomain linker strongly weakened binding to RNA, and concomitantly negatively impacted the degree of NP oligomerization (**Figure 4B, lanes 8-10** *versus* **11-13)**, further highlighting the importance of the NP interdomain linker for RNA binding, in addition to its contribution to HSPG binding on the cell surface **(Figure 3F, right panel)**. When the isolated NTD domain was tested for its RNA association properties by EMSA, we could not visualize any mobility shift, in contrast to CTD, for which clear shifts and oligomerization were evident (**Figure 4C, lanes 2-6** *versus* **7-11)**. Mixing individual domains together (**Figure 4C**, NTD+CTD) reveals stronger association with RNA than for CTD alone at the same concentration, suggesting that NTD has a positive effect on RNA binding in the presence of CTD, even if the two domains are not connected by a covalent linker, suggesting either cooperativity or an allosteric effect that enables a more productive binding of the RNA by the CTD (**Figure 4C, lanes 9** *versus* **12, 8** *versus* **13,** and **7** *versus* **15,** as indicated by bold arrows).

**Figure 4:**
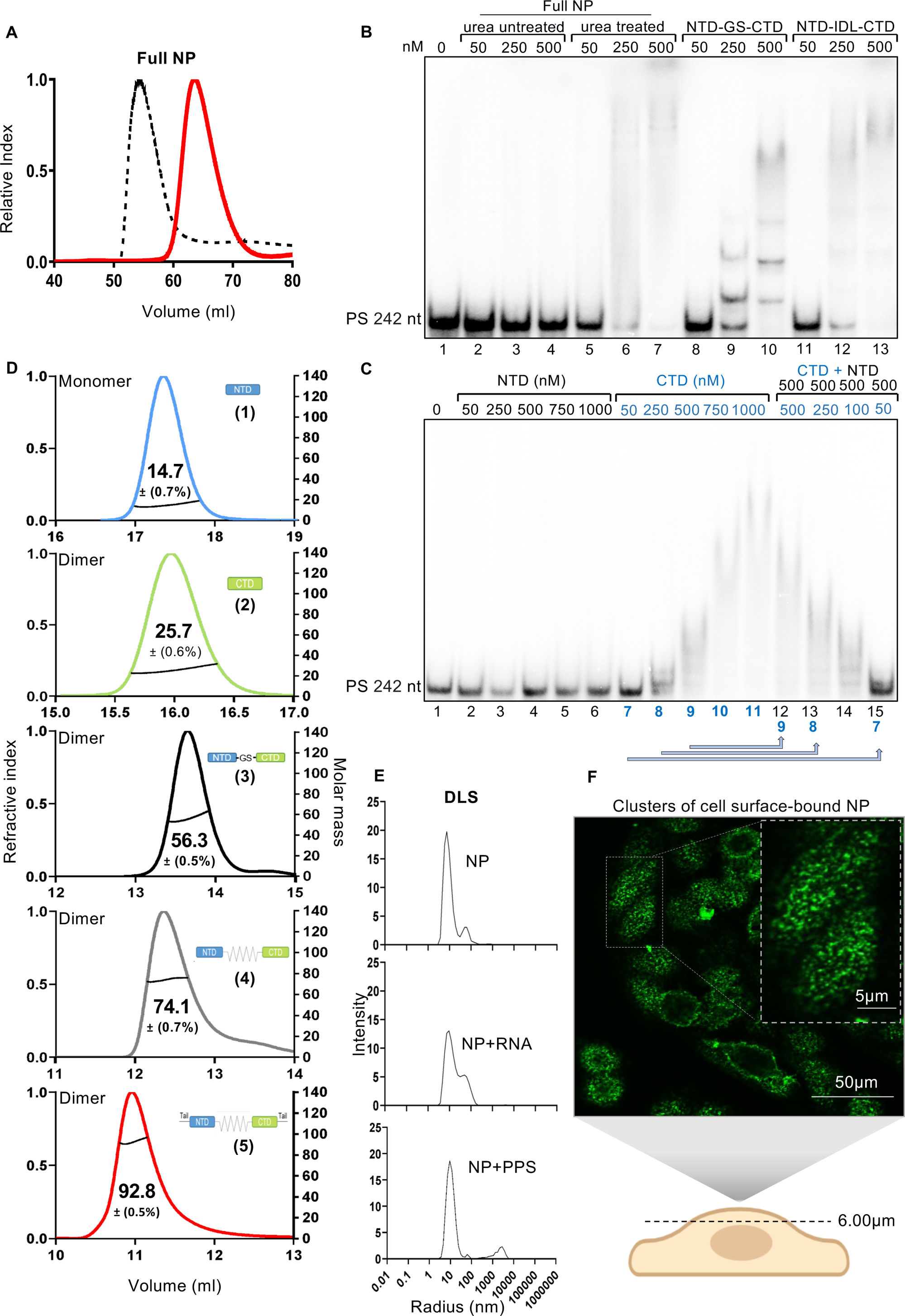
Binding of NP to RNA. **A:** Size exclusion chromatography (SEC) of wt full length NP with and without bacterial nucleic acid co-purifying contaminants. A Superdex (SD) 200 16/600 column was used for SEC. The black dashed line represents the elution profile of raw full-length NP prior to partially denaturing IMAC used to remove nucleic acid contaminants (see **Methods**), while the solid red line represents the elution profile of full-length NP after removal of co-purifying bacterial nucleic acids. The elution shift toward later elution volume indicates dissociation of oligomers upon removal of RNA. **B,C:** Electrophoretic mobility shift assay (EMSA) of end-labeled RNA (see **Methods** for details) and different NP constructs, as defined in Figure 3D. **B:** Labeled RNA was incubated with wt full length NP before and after removal of bacterial nucleic acid contaminants (lanes 2-4 and 5-7, respectively), NTD-GS-CTD (lanes 8-10), and NTD-IDL-CTD (lanes 11-13), at indicated concentrations. **C:** RNA was incubated with NTD (lanes 2-6), CTD (lanes 7-11), and a mixture of NTD + CTD (lanes 12-15). **D:** SEC coupled with multi-angle laser light scattering (SEC-MALS) analysis of NP constructs. Normalized values of refractive index (left Y axis) and measured molar mass in solution (right Y axis, Astra software, see Methods) are plotted against elution volumes. The line under each peak corresponds to the average molecular weight (MW) in solution (indicated inside the peak in kDa), as determined by MALS. Dimer/monomer at the left top corner of each graph indicates the apparent equilibrium monomer/dimer state concluded from the MW analysis versus expected MW of the monomer. **E:** Dynamic light scattering (DLS) shows changes in the average distribution of molecular radius as evaluated from scattering for wt full length NP only (top panel) or wt full length NP after incubation with RNA (middle panel) or with PPS (bottom panel), for 15 min at room temperature. **F:** Confocal fluorescence microscopy of A549 cells that were incubated with FITC-labeled NP (0.5μM). Live cells were labeled at 4°C to prevent endocytosis and then fixed prior to confocal imaging (see **Methods**). Note cell surface patches, indicating clustered distribution of surface bound NP. Scale bars of 50μm (main image) and of 5μm (inset) are shown. Schematic illustration at the bottom represents the shown focal plane, corresponding to the top of the cells magnified in the inset. Further optical Z-sections of the presented field are shown in **Supplementary Figure S3**.

To determine the oligomerization state and molecular weight of NP and the NP derived constructs in solution (in the ‘apo’ forms, free of RNA), we applied SEC-MALS (size exclusion chromatography coupled with multi-angle laser light-scattering, **Figure 4D**). While NTD is monomeric, CTD and all the NP derived constructs containing CTD form dimers in solution in the absence of RNA. We therefore hypothesize that a strong avidity contribution is responsible for the tighter RNA association of CTD and of CTD-containing NP derivatives, including the full-length NP. Of note however, our FACS experiments point out that for cell surface binding, no significant difference in affinity between the isolated CTD and NTD domains is observed; both domains, covalently linked together, are needed to achieve stable association (see **Discussion**). Dynamic Light Scattering (DLS) confirmed the tendency of NP to form high order oligomers in the presence of either RNA or the soluble heparin mimetics PPS (**Figure 4E**). Furthermore, inspecting uninfected A549 cells with surface bound FITC-labeled NP by direct fluorescence microscopy reveals a patched (non-homogeneous) distribution (**Figure 4F**, >0.2µm, defined by the Abbe diffraction limit), suggesting that NP forms macroscopic clusters upon surface binding, most likely reflecting the tendency of NP to oligomerize upon association with HSPGs. The latter observation is in line with the results of RNA driven oligomerization of NP, revealed by EMSA and DLS.

To summarize, these experiments support our model that NP attaches to cells by binding to sulfated proteoglycans, using its RNA binding sites. Moreover, both NTD and CTD are involved in binding, and in contrast to previous reports, it is the CTD domain that associates stronger with RNA and with heparin. Both domains, NTD and CTD are essential to achieve stable cell surface binding, which is significantly accentuated by the native interdomain linker. NP oligomerizes upon its association with RNA and with soluble heparin analogs, and by a similar mechanism forms large scale oligomers upon its binding to the cell surface of A549 cells. CTD is essential for oligomer formation, most likely through its dimerization domain.

Our model suggests that RNA and heparin binding sites on NP overlap. Analysis of the NP sequence conservation pattern highlights the conservation of these RNA/heparin binding sites, compared to the lower sequence conservation around and on the opposite side of the NTD and CTD domains (**Figure 3A**). Consequently, we can assume that it is the variable part of NP that remains exposed to recognition by antibodies, while the more conserved, basic RNA/HSPG binding patches are masked upon binding of NP to the cell surface.

### Recognition of surface-bound NP by anti-NP antibodies found in COVID-19 patient fluids

What are the possible immunological consequences of NP binding to uninfected cells? To answer this question, we tested whether human sera from recovered patients (anti-NP IgG^+^) recognize and bind NP attached to the cell surface. To this aim, we applied two complementary approaches: indirect immunofluorescence microscopy and cell-ELISA. We first used immunofluorescence microscopy to visualize live A549 cells with surface bound NP, using sera from COVID-19 recovered individuals (compared to naïve sera), followed by fluorescent secondary antibodies against human IgG. Strong cell surface staining was visible for A549 cells coated with NP and incubated with patients derived anti-NP positive serum, while no fluorescent signal was detected when NP was omitted, or when serum samples from naïve individuals were used, confirming the specificity of the immunostaining (**Figure 5A**). We next employed cell-surface ELISA for the quantitative evaluation of immunoreactivity toward surface bound NP by multiple clinical sera from the pre-vaccination SARS-CoV-2 epoch. We found a significant increase in reactivity when comparing sera from recovered to naïve individuals (n=54 and n=18, respectively, **Figure 5B**, left panel, *P value*<0.0001). Control experiments in which NP was omitted demonstrated the specificity of anti-NP IgG cell-surface ELISA based screening (**Supplementary Figure S4A**). Similar results were also observed for analysis of BAL fluids of recovered *versus* naïve patients (**Supplementary Figure S4B**).

**Figure 5:**
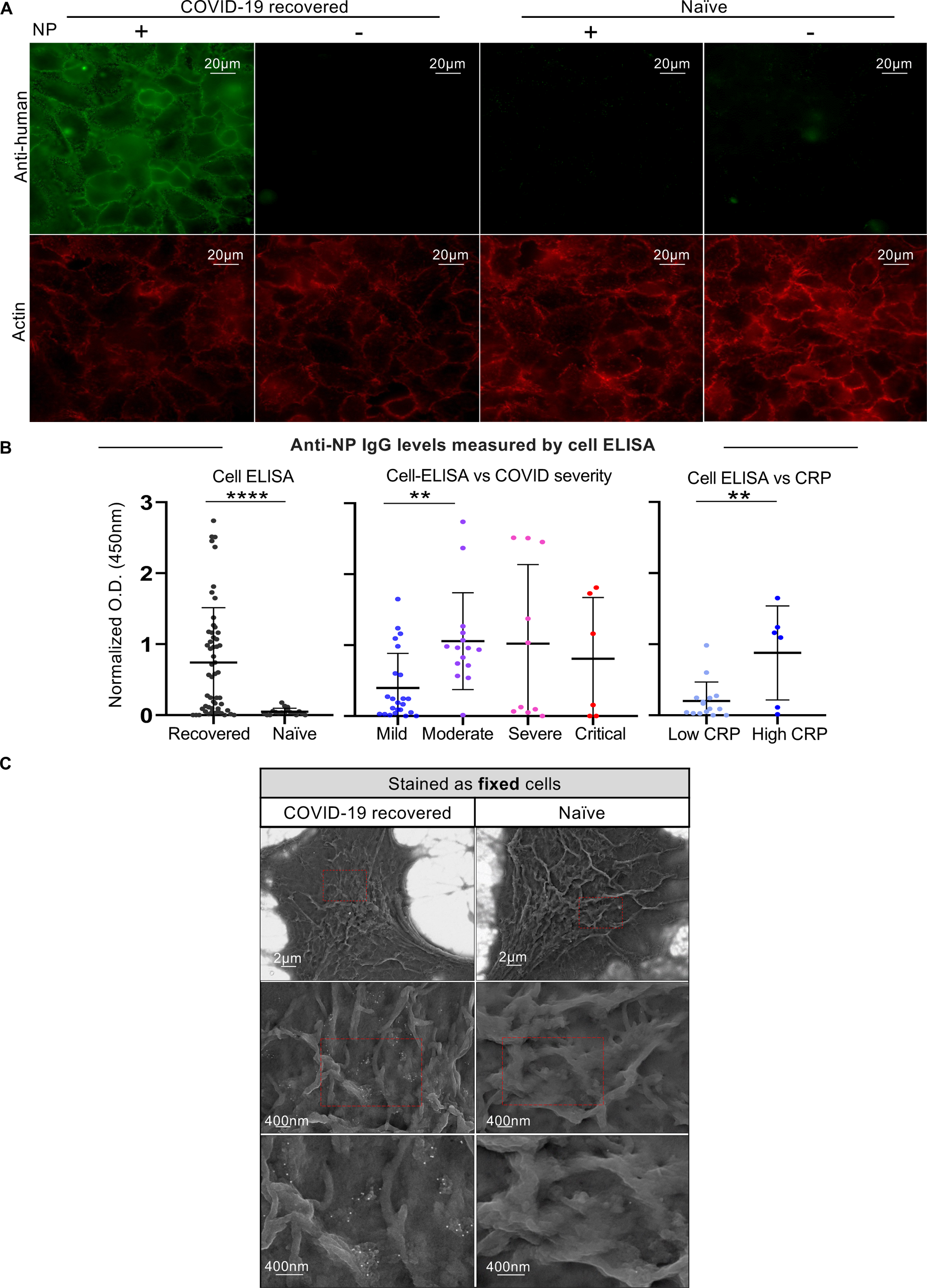
NP bound to the surface of uninfected A549 is recognized by antibodies from COVID-19 recovered patients. **A:** Immunofluorescence microscopy of A549 cells that were incubated with NP (0.65μM), washed, and subsequently exposed to plasma samples collected from either naïve or COVID-19 recovered individuals, followed by Alexa488 conjugated anti-human IgG secondary antibodies (upper panel). +/− signs above the images indicate the addition of NP (+) or its omission (-) upon staining of A549 cells. Cell outlines were revealed by phalloidin staining of F-actin filaments (lower panel). Scale bars represent 20µm, as indicated. **B:** Anti-NP cell-ELISA of A549 cells pre-incubated with full length NP at 4°C, and subsequently incubated with sera samples collected from naïve (n=18) or COVID-19 recovered individuals (n=60) (left panel). Middle panel: Corresponding results categorized by COVID-19 severity (mild, moderate, severe and critical, see **Methods**). Right panel: Mild cases were further sub-categorized based on the respective CRP levels (low CRP <7.5 *versus* high CRP >7.5). Student’s two-tailed unpaired t-test was used to compare recovered *versus* naïve (left panel), mild *versus* moderate (middle panel), and high CRP *versus* low CRP (right panel). *’**’* indicates *P value<*0.01, ‘****’ indicates *P value<*0.0001. The corresponding background staining values were subtracted from each ELISA measurement (see **Supplementary Figure S4A** for details). **C:** Immuno-gold Ultra-High Resolution Scanning Electron Microscopy (UHR-SEM) of A549 cells that were pre-incubated with full-length NP at 4°C and subsequently incubated with plasma samples collected from naïve or COVID-19 recovered individuals. Samples were then stained by goat anti human IgG secondary antibodies conjugated to 18nm colloidal gold particles for immunogold detection by SEM. The secondary staining was performed on fixed samples to avoid clustering artifacts of secondary immunostaining. Scale bars represent 2µm and 400nm, as indicated.

A key feature of cell-ELISA is that it detects solely anti-NP IgGs that target NP antigenic sites that remain exposed when the latter is bound to the cell surface (**Supplementary Figure S4C**). This is in contrast to classical ELISA (as well as the ARCHITECT anti-NP IgG test) that aim to detect all IgGs targeting NP. Despite these differences, the two assays are overall well correlated, indicating a general immunological dominance of epitopes displayed by cell surface-bound NP (Pearson correlation coefficient *r* = 0.86, see **Supplementary Figure S4D**). The relative magnitude of the two measurements, however, varied among the patients, suggesting patient specific paratope/epitope variations toward NP antigenic sites.

### Association between anti-NP IgG levels measured by cell ELISA and COVID-19 clinical state

In order to explore possible immunological correlates of COVID-19 clinical stages with anti-NP cell-ELISA values of the corresponding patients’ sera, we turned to a cohort of SARS-CoV-2 patients, for whom we had a collection of sera and the retrospective clinical snapshot data. The patients were stratified as mild, moderate, severe and critical cases, according to clinical information available, see **Supplementary Table S1** for classification, and **Methods**). Analysis of anti-NP cell ELISA values reveals a significant difference between the mild and the moderate cases categories of COVID-19 patients (**Figure 5B**, middle panel; *P value*=0.0013). The main clinical difference between these two groups is the presence of lung radiological abnormalities in moderate *versus* mild cases, as determined by chest X-rays and/or computerized tomography scans (see **Methods**). This observation suggests a correlative link between lung pathologic states and the levels of IgG antibodies in circulation that recognize cell surface bound NP. Of note, the corresponding differences of either total anti-NP IgG levels (as measured by ARCHITECT anti-NP IgG test) or total anti-RBD IgG levels (as measured by custom laboratory ELISA) are less significant (**Supplementary Figure S4E-H,** see also **S4C** and **Discussion**).

Closer inspection of the mild cases reveals a small subset of patients with high values for anti-NP cell-ELISA IgGs (**Figure 5B**, middle panel). These could be cases that are about to deteriorate, but do not yet show lung pathological changes when examined by chest radiology. Since our cohort study contains only snapshot clinical and radiological information at the time of sample collection and hospital admission, we decided to characterize the mild cases further using C-reactive protein (CRP) values available from blood tests. CRP is usually produced by liver cells, rising upon inflammation and infection, especially during the acute stage, and was found to correlate with severity of COVID-19 (3). Indeed, for the mild cases, sera samples from high CRP patients (>7.5mg/dl) exhibited significantly higher anti-NP reactivity in cell-ELISA compared to sera from patients with low CRP (<7.5mg/dl) (**Figure 5B**, right panel, *P value*=0.0029) whereas the levels of total anti-NP IgG (ARCHITECT anti-NP IgG test) between these two groups were not significantly different, see **Supplementary Figure S4E,G**).

Overall, these results suggest an association between high titers of anti-NP IgG recognizing exposed epitopes on the cell surface bound NP and between more severe COVID-19 clinical cases, as defined either by (i) the presence of lung radiological abnormalities (*i.e.*, mild versus moderate cases), or by (ii) high CRP levels (among the mild cases). We next focus on characterizing the immunological mechanism that could explain this association.

### Anti-NP IgGs activate the classical complement pathway, targeting uninfected cells marked by surface bound NP

In order to further explore an immunological skew scenario of biased attack of uninfected cells, we asked whether the immune system, driven by Fc effector functions, could target NP-marked cells in individuals with anti-NP IgG. In the current study we focused on the complement system in light of the reported contribution of the complement to the pathophysiology of COVID-19 (35–37). Complement recognition of antigenic complexes is known to require several threshold prerequisites, one of which is high degree of clustering of the antigenic sites (15,38). In this view, we observed that NPs form clusters on the cell surface, as revealed by puncta appearance of cell surface bound fluorescent NP (**Figure 4F**) and by NP-IgG surface complexes (**Figure 5A**), in agreement with the tendency of NP to oligomerize in association with RNA or PPS (**Figure 4E**).

To further explore the ultrastructure of NP-antibody immune complexes on the surface of uninfected cells after incubation with anti-NP positive sera from recovered patients, we used immunogold ultra-high resolution scanning electron microscopy (UHR-SEM, **Figure 5C**). The immunogold clusters visualized by UHR-SEM reveal the distribution of anti-NP human IgGs on the cell surface of uninfected cells with NP bound to their surface. Importantly, immunogold clusters were only visible after addition of NP^+^ antisera from COVID-19 recovered patients, but not of naïve sera (**Figure 5C**). Furthermore, no signal was obtained when the NP binding step was omitted, demonstrating the specificity of immunogold staining (**Supplementary Figure S4I**). Of note, to avoid artificial non-physiological clustering events by secondary polyclonal antibodies used for detection in immunogold UHR-SEM, we introduced a fixation step prior to the secondary immunogold detection (see **Methods**).

Our observation that NP-antibody complexes clustered on the surface of uninfected cells suggests that these complexes might provide a target for the classical complement pathway, attacking large antigenic assemblies (15). To mimic such a situation *in vitro*, we used A549 cells coated with recombinant NP, and subsequently incubated these cells with sera from patients with anti-NP IgG, followed by incubation with guinea pig derived complement. Sera of patients and of naïve control sera were heat inactivated to eliminate the endogenous complement activity before adding guinea pig complement for the assay, in order to avoid possible variations in the endogenous complement state. The complement activation/deposition was monitored by immunostaining of the cells with fluorescent anti-C3b antibodies that specifically recognizes guinea pig C3b. C3b is a product of C3 convertase activity, providing a marker for complement activation. Analysis by immunofluorescence microscopy revealed C3b complement deposition on A549 cells, exclusively by sera of patients with anti-NP IgG, and only in the presence of NP (**Figure 6A**). This finding demonstrates the activation of the classical complement pathway mediated by exogenous surface bound NP and by anti-NP antibodies found in serum samples from COVID-19 recovered individuals. We next performed FACS on a set of human sera from COVID-19 recovered individuals to compare the IgG reactivity in cell ELISA toward cell surface NP versus amount of C3b complement deposition (**Figure 6B**, left and right, respectively). The results suggest that while presence of IgGs against cell surface bound NP is a prerequisite, additional factors are involved in determining the amount of C3b deposition. For example, while samples P1 and P2 display almost identical anti-NP IgG levels in cell surface FACS and both samples induce significant C3b deposition, they exhibit a strong difference in the specific amounts of C3b deposition (**Figure 6B**, compare the two upper panels, left and right). These differences may be explained either by IgG subclass repertoire or by sub-variations in targeted epitopes, and are further addressed in the **Discussion**.

**Figure 6:**
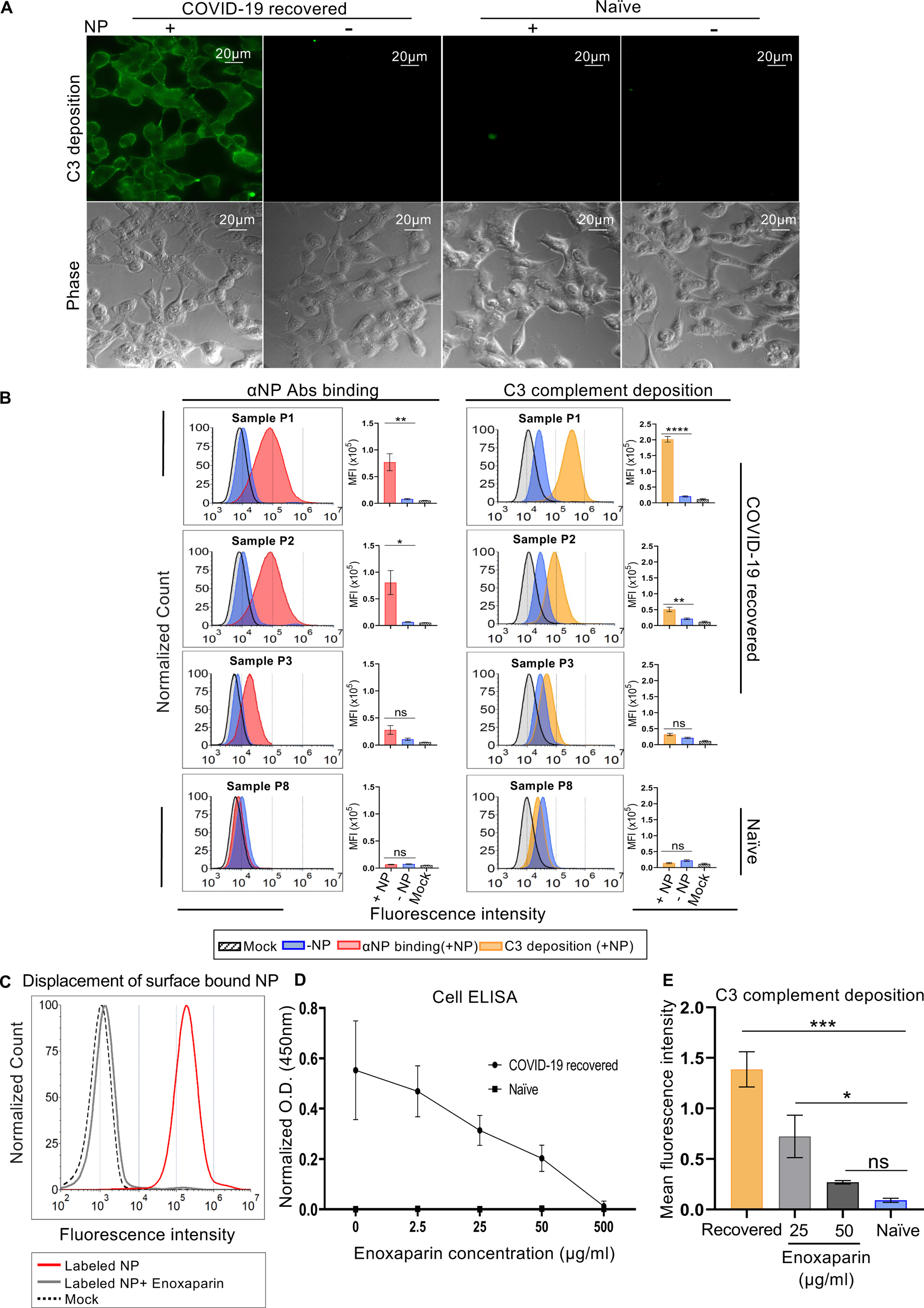
Anti-NP antibodies from plasma of COVID-19 recovered patients bind to cell surface-bound NP and induce C3b complement deposition. **A,B:** Cell-surface bound NP activates complement deposition, shown by immunofluorescence microscopy and FACS**. A:** Immunofluorescence microscopy of A549 cells that were either incubated (+) or not (-) with NP and subsequently exposed to heat-inactivated plasma samples collected from either COVID-19 recovered or naïve individuals, followed by exposure to heterologous complement from Guinea pig serum. C3b complement deposition was revealed by FITC conjugated goat anti-Guinea pig C3b. Scale bar represents 20µm. **B:** Corresponding flow cytometry experiment for four different plasma samples collected from either COVID-19 recovered or naïve individuals (P1-P3 and P8, respectively). To compare complement C3b deposition to anti-cell surface bound NP IgG reactivity, samples were incubated as above for detection of C3b complement deposition (**right panel**), or with donkey anti-human IgG, secondary antibodies conjugated to Alexa488 for detection of total IgG reacting with cell surface bound NP **(left panel)**. In all experiments, 0.65μM full length NP was added, and flow cytometry histogram plots show the mean fluorescence intensity values of three independent replicates (mean +/− SEM).**C-E:** Enoxaparin prevents NP-induced complement activation. **C:** Flow cytometry of live HeLa cells that were incubated at 4°C either with FlTC-labeled full length NP, or co-incubated with FITC-labeled full length NP and enoxaparin (500μg/ml). **D:** Anti-NP IgG Cell-ELISA of A549 cells in the presence of enoxaparin. A549 cells were incubated at 4°C first either with full length NP or co-incubated with full length NP and enoxaparin applied at different concentrations, as indicated. Cells were subsequently incubated with plasma samples collected from COVID-19 recovered or naïve individuals and next with donkey anti-human IgG conjugated to HRP. The signal was developed using TMB substrate (normalized OD 450nm values are presented). **E:** Flow cytometry of complement deposition (C3b) mediated by anti-NP antibodies in the presence of enoxaparin. A549 cells were incubated at 4°C with either full length NP only, or co-incubated with full length NP and enoxaparin (25μg/ml and 50μg/ml). Cells were subsequently incubated with heat-inactivated plasma samples collected from COVID-19 recovered or naïve individuals. Next cells were incubated with Guinea pig complement source and deposition of C3b was evaluated by goat anti-Guinea pig C3b conjugated to FITC (see **Methods**). One-way analysis of variance (ANOVA) and Tukey’s multiple comparisons test were used to compare all conditions in flow cytometry histogram analysis. ‘ns’ (non significant) indicates *P value* >0.05; ‘**’ indicates *P value* < 0.01; ‘***’ indicates *P value* <0.001; and ‘****’ indicates *P value <*0.0001.

The electrostatic nature of interactions between NP and the sulfated proteoglycans on the cell surface, together with the capacity of the sulfated soluble heparin analog, PPS to displace cell surface bound NP (**Figures 2F,G**) suggest promising novel therapeutic opportunities to interfere with NP mediated complement targeting of uninfected cells. In order to explore this possibility, we asked whether incubation with enoxaparin, a widely used low molecular weight heparin-based anticoagulant, could displace NP and thereby rescue uninfected NP-coated A549 cells from anti-NP IgG mediated complement deposition. As expected, enoxaparin was able to displace fluorescently labeled NP, as revealed by FACS (**Figure 6C**). Moreover, enoxaparin showed a dose dependent capacity to reduce cell surface binding of anti-NP IgGs (**Figure 6D)**, in line with the observed NP displacement (**Figure 6C**). Accordingly, enoxaparin exhibited potent rescue of A549 cells from anti-NP IgG driven complement deposition (**Figure 6E**, see also **Supplementary Figure S5**), revealing a novel, previously not yet considered mechanistic facet of immunological COVID-19 management with soluble heparin analogs, in addition to their previously reported benefits such as inhibition of SARS-CoV-2 entry (39,40) and prevention of blood clotting (41).

To summarize, in this study we demonstrate that NP of SARS-CoV-2 plays an important role beyond its binding and packaging of viral RNA. We show that NP uses its conserved RNA binding sites to also attach to the HSPG layer on the surface of uninfected cells (**Figure 7**, (**1**)), exposing a less conserved surface to the immune system (**Figure 7**, (**2**)). Subsequent binding of anti-NP IgGs from patients sera triggers the deposition of complement (**Figure 7**, (**3**)), targeting these NP coated, uninfected cells and leading to tissue damage. This can be prevented by a soluble heparin analog, enoxaparin (**Figure 7, bottom**). Our findings reveal a mechanistic link between several previous clinical and diagnostic observations related to the association of COVID-19 severity with increased titers of anti-NP antibodies (7,8) and systemic complement activation (35,36,56,57). Moreover, we substantiate the connection to observed lung tissue damage in patients with COVID-19 complications, by detecting NP targeting IgG response in the BAL fluid collected from recovered patients that underwent bronchoscopy.

**Figure 7.**
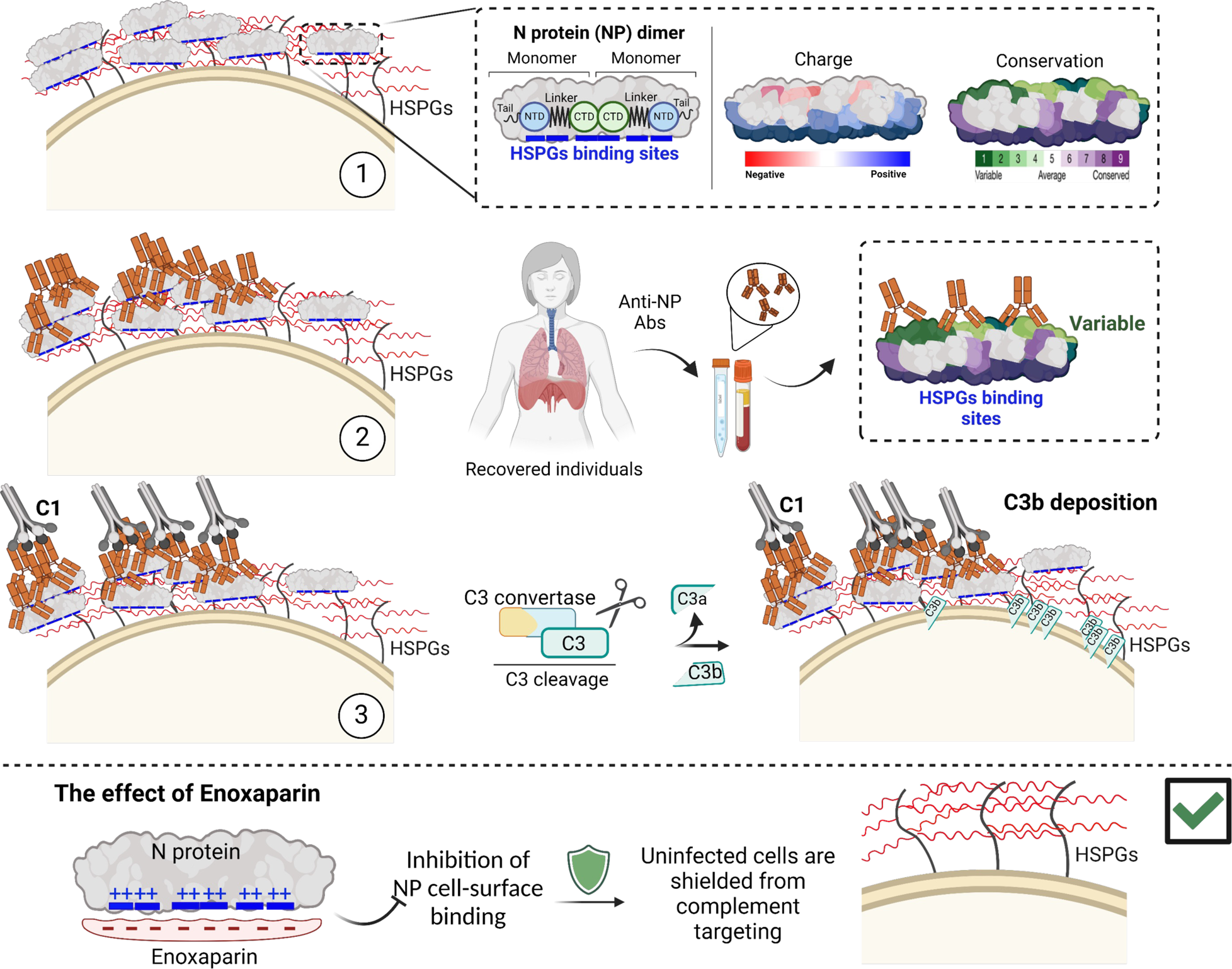
Model of the suggested functional impacts of extracellular NP and the associated immune response mechanisms. See text for more details.

## Discussion

Traditionally, NP has been described as an intraviral or intracellular protein, based on its major function of binding to viral RNA and facilitating its packaging within the virion (42–44). Here we show that NP plays additional roles in the extracellular medium. We report that anti-NP IgGs in the serum of COVID-19 recovered individuals (in ELISA screens of sera samples collected from individuals who recovered in the pre-vaccination era) recognize NP-covered uninfected cells. This not only reconfirms that NP is exposed to the immune system (7,8,21), as shown by recent reports of NP secretion to the surface of infected cells and its subsequent transport to by-standing neighbors (10,11),but more importantly indicates that NP-bound healthy tissue in proximity to the site of infection might be targeted by anti-NP antibodies and destroyed by complement deposition, thereby diverting in part the immune response from infected to healthy cells. This can explain the significant damage to uninfected tissues seen in COVID-19 patients (2).Cell surface clusters of antigen-antibody complexes serve as an optimal substrate for the binding of the C1q component, ultimately initiating complement system activation (38,45). Large scale clusters of cell surface bound NP, as well as of polyclonal anti-NP IgG immune complexes visualized as cell surface assemblies, most likely reflect extended HSPGs with their repetitive linear sulfated GAGs, providing polymeric NP binding sites, instrumental in alarming immune responses.

Using samples from a well characterized SARS-CoV-2 early pre-vaccination cohort, which was classified based on lung radiological abnormalities (chest X-rays and/or computerized tomography scans), we show significant association of lung damage with higher titers of anti-NP antibodies capable of binding the surface of A549 cells coated with recombinant NP. Moreover, BAL samples from recovered individuals also displayed reactivity toward cell surface bound NP, suggesting that complement activation could indeed account for alveolar pathophysiology of COVID-19. To fully explain the extent of complement activation, it will however be necessary to consider additional factors, since our results show that similar IgG reactivity can still result in different degrees of complement activation. This may include the Fc determinants of anti-NP IgGs, IgG subclass and glycosylation which have been reported to play a significant role in mediating a plethora of Fc mediated functions, such as complement deposition (15,46). Moreover, it has been reported that NP can aggravate inflammation by a serine-protease-mediated lectin complement pathway overactivation through an IgG independent pathway (47).

Presence of cell surface bound NP was reported not only for SARS-CoV-2 and other human cold coronaviruses (10,11), but also for NP of Influenza A virus infected P815 mast cell line (49). It is not yet clear whether NPs of other viral families also follow a similar fate of cell surface translocation and transmission to neighbors, and if so, whether they elicit antibody responses that may target bystanders and cause associated secondary tissue damage. While the exact mechanism of secretion and bilayer translocation is not yet deciphered, additional cellular cytosolic RNA binding or nuclear RNA/DNA binding proteins like Ro/SSA and La/SSB proteins were reported to be present on the surface of cells in health (50) and disease (50), upon viral infections (51) and in particular, upon severe autoimmune disease conditions like Sjorgen, Lupus and multisystem inflammatory syndrome in children (MIS-C) (52,53). Of note, recent work identified previously unknown small RNA molecules presented on the surface of mammalian cells through N-linked glycans (54,55). Whether these small RNA molecules are engaged in interaction with, or contribute to a translocation of viral or cellular RNA binding proteins remains yet to be investigated.

Investigation of the interaction of NP with heparin, cell surface, and RNA reveals that while these substrates bind isolated NTD and CTD, their association is much stronger in the context of full-length NP and that the linker plays a critical role. Comparison of structural models of an NP dimer bound to heparin and to RNA reveals that in both cases the two NTD domains and the CTD dimerization domain bind to functional units of the substrate, but that their relative orientation may differ significantly, adopting an extended conformation when bound to linear sulfated GAGs, but a wrapped compact conformation around RNA (20). These models highlight the contribution of the flexible, positively charged linker: Thanks to its flexibility, it enables the distinct orientations between NTDs and CTD, and thanks to its positive charge, it can by itself contribute an additional site for binding to the negatively charged substrates.

Tighter association of NP with RNA was previously reported for a construct spanning NTD-linker-CTD compared to the isolated domains added together, using fluorescence polarization (48). Moreover, Hydrogen Deuterium exchange (HDX) experiments showed that linker accessibility is significantly changed upon binding of the NTD to RNA, supporting a possible additional direct interacting site between the linker and RNA (48), in agreement with structural models that suggest its involvement in binding RNA (20). Thus, while overall our findings on RNA binding modes are along the lines with other recent publications, the specific contributions of the different parts of NP to cell surface binding have not been previously explored.

Despite similarities in binding determinants to surface HSPGs and RNA, our experimental results show also specific differences. Regarding RNA binding, EMSA experiments detected significant binding and oligomer formation only for isolated CTD, but not NTD. In turn, NTD substantially increased binding of CTD to RNA, even when both domains were added separately, pointing to an allosteric mechanism. Thus, while crystal structures of NTD-RNA complexes have been reported, no such structure has been solved yet for CTD-RNA, full-NP-RNA, CTD-Heparin or full-NP-Heparin, most likely due to challenges posed by oligomer formation of the latter. Regarding the association with sulfated GAGs, individual domains bound stably to the heparin column at low ionic strength, while we had to omit the washing step in order to observe their binding to the cell surface at isotonic conditions, indicating a high off-rate of single NTD and CTD binding to the cell surface. Regarding the role played by the N- and C-terminal tails, they significantly reduce cell surface binding, but did not elicit any observable effect upon binding to heparin column or to RNA. Overall these results highlight differences of NP association with cell surface linear sulfated GAG chains in comparison to the smaller heparin and short RNA fragments.

The results of the current study show that enoxaparin effectively hinders the binding of NP to the cell surface in a dose-dependent manner and as a result significantly reduces the levels of C3b deposition, exposing another, yet unreported benefit of heparin analogs beneficial usage for COVID-19 management in addition to their known roles in clot prevention and inhibition of spike mediated virus entry (39–41).

A common property of viral NPs and cellular RNA/DNA binding proteins is their electrostatic/basic regions, engaged in interaction with nucleic acids. In the current work we analyze structural details and constraints of geometric arrangements upon interaction with RNA versus cell surface HSPGs. Furthermore, findings presented in the current study show that soluble low molecular weight heparin analogs like enoxaparin, a widely used anticoagulant, dissociates SARS-CoV-2 NP and rescues uninfected cells from complement mediated attack. This raises the possibility that similar strategies may apply to alleviate exacerbated immune pathologies, associated with additional viral diseases and relieve certain autoimmune damaging conditions like Sjorgen, Lupus and multisystem inflammatory syndrome diseases.

## Materials and Methods

### Cloning and mutagenesis

The NP constructs NP_41-174_ (corresponding to NTD in the main text), NP_246-364_ (corresponding to CTD in the main text), NP_41-364_ (corresponding to NTD-IDL-CTD in the main text) and full-length NP were generated using Gibson assembly (Gibson assembly master mix, New England Biolabs) according to the manufacturer’s protocol and are cloned in pET15b.

The mutation of NTD_GS_CTD was generated by PCR and Gibson assembly. A list of all primers is provided in **Supplementary Table S3**. All the constructs were verified by DNA sequencing.

### Protein expression and purification

NP constructs were expressed in *E. coli* T7 express (New England Biolabs) cells. The transformed cells were grown in 2xYT and induced at OD (A600nm) of 0.6-0.8, at 16°C overnight with 0.3mM isopropyl-β-thio-galactoside (IPTG). Subsequently, cells were harvested via centrifugation at 6,000 x g for 15 min and stored at −80°C. Cell pellets containing the expressed NP constructs were resuspended in lysis buffer (50mM NaH_2_PO_4_, pH 7.0, 500mM NaCl, 3M Urea, and 5mM β-mercaptoethanol), supplemented with 1 mM phenyl-methyl sulphonyl fluoride (PMSF) and DNase I. Disruption was achieved using a microfluidizer (Microfluidics). The lysate was clarified by centrifugation at 47,000 x g for 45min and applied to 5 ml His-Trap columns (GE Healthcare). After washing with a buffer excluding urea, elution was performed with a linear imidazole gradient of 10–250mM in 30 column volumes (CV). Purified protein fractions were pooled and dialyzed overnight at 4°C against dialysis buffer (20mM NaH_2_PO_4_, pH 7.0, 500mM NaCl, and 5mM β-mercaptoethanol) in the presence of His-tagged TEV protease. The cleaved protein underwent a second round of purification via His-Trap column, and the flow-through fraction containing the cleaved protein was collected. Further purification was accomplished using a 16/600 Superdex 75 pg size-exclusion chromatography column (GE Healthcare), equilibrated in protein buffer (20mM Tris, pH 7.8, 150mM NaCl). All proteins were concentrated, flash-frozen in aliquots in liquid N_2_, and stored at −80°C.

### Crystallization

The CTD crystals were grown at 20°C using the hanging drop vapor diffusion method. The protein was concentrated to 13 mg/ml and crystallized in a solution containing 200mM NaCl, 100mM sodium/potassium phosphate at pH 6.5, and 25% PEG1,000. After 72 hours, the crystals were cryoprotected using a reservoir solution containing 25% glycerol and then cryo-cooled in liquid nitrogen.

### Data processing and structure refinement

The crystal diffraction data were collected by the rotation method on beamline ID23-2 at ESRF synchrotron at 100K. Data was processed by XDS (58). The structure was solved by the molecular replacement method implemented in the MoRDa pipeline (59). The best model selected by MoRDa was a monomer of PDB ID 6WZO (18). Further structural studies were carried out using the CCP4 program package (60) in the CCP4CLOUD interface (61). The structure was refined by REFMAC5 (62) and rebuilt in COOT (63). Details of the quality of the refined model are presented in **Supplementary Table S2**.

### Protein labeling

NP was labeled with Fluorescein isothiocyanate (FITC) using a sortase recognition peptide N-terminally conjugated to FITC (FITC-LPETGG). NP was incubated with this peptide and the sortase A enzyme (Sortase A pentamutant (eSrtA), Addgene #75144) (64) for 1.5 hours at 25°C in the reaction buffer (50mM Tris pH 7.5, 150mM NaCl, 10mM CaCl_2_). Sortase A is a transpeptidase from *Staphylococcus aureus* that cleaves the Thr-Gly bond at the recognition site LPXTG and forms an acyl intermediate with Thr (65), which subsequently reacted with the N-terminal Gly of NP, resulting in the FITC labeled peptide fused to the N terminus of NP (see **Figure 2A**). Subsequently, NP was purified using Sephadex G-25 columns.

### Binding to heparin column

NP constructs (each at 100 μg) were loaded separately on 1 ml heparin column (GE17-0407-01, GE Healthcare) pre-equilibrated with buffer A (20mM Tris pH 7.5, 100mM NaCl). The column was then washed with 5 CV of buffer A and protein was eluted using a linear 10 CV gradient of 100mM to 1M NaCl.

### Dynamic light scattering (DLS) analysis

The mean hydrodynamic diameter determination of NP with and without PPS and RNA was performed using a Möbius instrument, at 540 nm laser wavelength, equipped with a 532 nm long pass filter (Wyatt Technology Corporation, Santa Barbara, CA 93117 USA). Each sample contained 50 µl of NP (2.7 mg/ml). Either PPS (5 µg) or RNA (1.85 µg) were added to the NP sample, mixed and incubated for 15 minutes at room temperature (RT). For each sample 5 runs were performed at 25°C.

### Western blot

Following electrophoresis, the proteins were transferred to a polyvinylidene difluoride membrane (110V, 1h, 4°C) using a Mini Trans-Blot cell apparatus (Bio-Rad). Non-specific binding sites were blocked using PBS/Tween 20 (0.05% Tween 20, J.T.Baker) containing 5% non-fat dry milk for 1h at RT. The membrane was then incubated with mouse anti-His antibody (catalog number #6200203, Bio-Rad), diluted 1:1,000 in the blocking solution and incubated overnight at 4°C, washed and overlaid with HRP-conjugated goat anti-mouse secondary antibodies diluted 1:5,000 in blocking solution for 1h at RT (Jackson ImmunoResearch, #115-035-003) and subsequently washed. Before imaging (using the ChemiDoc MP imaging system, Bio-Rad), the membrane was incubated in EZ-ECL solution (Biological Industries) for 5 min.

### Size exclusion chromatography with multi-angle light scattering (SEC-MALS)

NP and its derivatives were loaded on an analytical SEC column (Superdex 200 10/300 GL) equilibrated with a buffer containing 20 mM Tris, pH 7.5, 500 mM NaCl, and 1 mM DTT. The molecular weight within the chromatographic peak was determined by MALS using ASTRA software, v7 (Wyatt Technologies).

### Clinical cohort

The serum and plasma cohort used in this study were obtained by collaboration with the Hadassah Clinical Laboratory and Hadassah Medical Center’s blood bank, respectively. The Institutional Review Board (IRB) approval number for the study and analysis of convalescent samples which were diagnosed by RT-PCR for SARS-CoV-2 infection at the Hadassah Clinical Virology Laboratory, is 0235-20-HMO. The pre-COVID-19 controls were randomly retrieved from the blood bank serum samples that had been routinely tested at the Clinical Virology laboratory between 06-09/2019. The samples were kept at −80°C. The clinical data of the sera cohort were reviewed systematically by a certified pulmonologist. Clinical data included COVID-19 severity, lab results (WBC, CRP, DDM, ferritin) and imaging (chest X-rays and/or computerized tomography scans) (**Supplementary Table S1D**). COVID-19 severity was classified according to the World Health Organization severity scale. The reviewer pulmonologist had no access to any of this study’s cell-ELISA and other experimental results.

The sub-cohort of BAL and serum samples used in this study were obtained as part of a previous study (66). This study was approved by the IRB (0400-21-HMO) and each participant had to sign an informed consent form. BAL was obtained during medically indicated bronchoscopy performed at the Institute of Pulmonary Medicine at Hadassah Medical Center in Jerusalem, according to internationally accepted guidelines. The samples (serum, plasma and BAL) were maintained at –80°C. The samples were heat inactivated prior to application in the different assays (ELISA, cell-ELISA, IF, FACS, complement C3b deposition).

### Cell lines

A549 (A549 - CCL-185, ATCC) and HeLa (HeLa - CCL-2 - ATCC) cells were grown in Dulbecco’s modified Eagle’s medium (DMEM) supplemented with 10% (v/v) Fetal Bovine Serum, penicillin (100 IU/ml), streptomycin sulfate (100 µg/ml), and L-glutamine (2 mM). The cells were maintained in a 37°C humid incubator under 5% CO_2_ and routinely tested for contamination by mycoplasma.

### Complement

Lyophilized guinea pig complement (Sigma-Aldrich, 234395-5ML), was resuspended according to the manufacturer’s instructions (5 ml of ultra-pure DNase/RNase free DDW and stored in aliquots at –80°C).

### Indirect Enzyme-Linked Immunosorbent Assay (ELISA)

Antigens (NP, NTD, RBD) were coated at 4°C overnight onto MaxiSorp^TM^, 96-well plates (Thermo) in antigen dilution buffer (20mM Tris pH 7.5, 50 mM NaCl). Plates were blocked with 3% fat UHT milk diluted with PBS (1:1) for 30 min at RT. Sera samples were heat-inactivated (60°C, 30 min), serially diluted in blocking buffer, added to the wells, and incubated for 30 min at RT. Wells were rinsed (3x PBS), incubated at RT (45 min) with HRP conjugated secondary donkey anti-human IgG (H+L) (1:5,000) (Jackson ImmunoResearch), thoroughly washed and developed with 3,3′,5,5′-Tetramethylbenzidine (TMB) substrate. The reaction was stopped with 0.1% H_2_SO_4_ and the OD at 450 nm was quantified using a plate reader (Spark, Tecan).

### Commercial anti-NP IgG detection assay

In addition to the classical ELISA assay, we also used the ARCHITECT i2000SR high-throughput system anti-nucleocapsid (N) IgG (Abbott, Illinois, USA), diagnostic kit.

### anti-NP cell-ELISA

1.5×10^4^ A549 cells were plated overnight in 96-well plates. The cells were chilled on ice and washed 3x using ice cold PBS (containing Ca^2+^ and Mg^2+^). All the subsequent incubations until substrate development were performed at 4°C. Cells were incubated for 30 min with NP (0.65μM), washed (3x with PBS) and incubated either with serum or BAL fluid samples, collected from naïve or recovered individuals, for 30 min. Serum and BAL samples were diluted 1:50 and 1:6, respectively, using 1% BSA solution in PBS (Ca^2+^ and Mg^2+^), ultra-centrifuged for 15 min, 120,000 g_AV_ at 4°C. Cells were washed (3x PBS) and incubated with HRP conjugated donkey anti-human IgG (H+L) secondary antibodies (1:5,000) for 30min, washed and developed with TMB substrate (5 min incubation at RT). The supernatant was moved to a clean 96-well plate that contained 0.1% stop solution (0.1% H_2_SO_4_) and the OD at 450 nm was quantified using a plate reader (Spark, Tecan). Cells were then fixed using 4% formaldehyde in PBS for 30 min at RT and stained using Amido Black [50% Isopropanol(v/v), 20% acetic acid (v/v), 0.2% Naphthalene Black (w/v)] for 30 minutes at RT. The wells were then washed 3x using destaining solution [50% Isopropanol (v/v), 20% acetic acid(v/v)] and the bound dye was eluted by 50 mM NaOH solution. The supernatant was collected and OD was measured at wavelength 405 nm and the values were used for normalization of cell quantities between the wells.

### Binding of FITC-labeled NP and fluorescence microscopy

2×10^4^ HeLa cells were plated overnight in an 8-well coverglass bottom chambered slide (ibidi, 80827-90). The slide was chilled on ice and washed (3x cold PBS, containing Ca^2+^ and Mg^2+^) and all the subsequent incubations until the fixation step were performed at 4°C. NP was diluted in 1% BSA solution in PBS to a concentration of 0.65 μM, then added to the relevant wells and incubated for 30 min. Finally, cells were washed 3 times with PBS and then fixed using 4% formaldehyde for 30 minutes at RT. Competition binding assays were performed at the same condition, but cells were incubated twice with NP: First with FITC-labeled NP and then with unlabeled NP protein/competitor. Images were acquired using either widefield fluorescence microscopy (Nikon microscope air objective, 20x/0.4 NA), or confocal fluorescence microscopy (Nikon Microscope equipped with Yokogawa W1 Spinning Disk, ORCA-Fusion BT camera, pinhole size 50 µm, air objective, 40x/0.95 NA).

### Anti-NP antibody cell-surface binding

4×10^4^ A549 cells were plated overnight in an 8-well coverglass bottom chambered slide (Ibidi). Cells were chilled on ice and washed (3x cold PBS, containing Ca^2+^ and Mg^2+^) and the subsequent steps were performed at 4°C. NP was diluted in 1% BSA solution in PBS to a concentration of 3.25 μM, added to the relevant wells, and incubated for 30 min. Cells were then washed and incubated with plasma samples collected from naïve or recovered individuals for 30 min. Plasma samples (collected in the K3EDTA tubes and subsequently separated from cells by centrifugation) were diluted 1:100 using 1% BSA, ultracentrifuged for 15 min, 120,000 g_AV_ at 4°C. Following incubation, wells were washed and incubated with Alexa488 conjugated donkey anti-human IgG (H+L) at a dilution of 1:200 for 30 min. Finally, cells were washed and fixed using 4% formaldehyde at RT for 30 minutes. Images were acquired using an epifluorescence microscope (Olympus IX83 microscope oil objective, either 60x/1.42 NA, or 100x/1.45 NA, as indicated) and processed using Fiji ImageJ software. A similar protocol was applied for the flow cytometry assay, for which 4×10^5^ A549 cells were plated per well in a 12-well plate overnight. Following the incubation with the secondary antibodies the cells were detached using 10 mM EDTA, washed and centrifuged at 500 x g for 5 min, and analyzed by Cytoflex flow cytometer. 30,000 events were recorded. The data analysis was conducted using FCS Express software (De Novo Software, Pasadena).

### Anti-NP antibody-dependent complement deposition (ADCD)

5×10^4^ A549 cells were plated overnight in an 8-well coverglass bottom chambered slide (ibidi) coated with 2.5 μg/cm^2^ Cell-TAK (Corning®, 354240). Cells were grown in DMEM, 0.5% FBS, 10% Pannexin NTS, a defined serum substitute (PAN-Biotech, Germany) prepared in conditioned medium. Conditioned medium was collected 48h after addition to A549 cell culture, centrifuged and filtered (0.45μm). The cells were chilled on ice and washed 3x with ice cold PBS (containing Ca^2+^ and Mg^2+^) and all the subsequent steps were performed at 4°C. Cells were next incubated for 30 min with NP (3.25μM), washed and incubated with plasma samples (diluted 1:20), as described above. This was followed by incubation with guinea pig complement (diluted 1:20 in veronal buffer) for 30min. Cells were then washed and incubated with FITC-conjugated goat anti-guinea pig complement C3 (dilution of 1:500), for 30 min and subsequently fixed using 4% formaldehyde for 30 minutes at RT. Images were acquired using epifluorescence microscopy (Olympus microscope oil objective (60x/1.42 NA, or 100x/1.45 NA, as indicated) and processed using Fiji ImageJ software. A similar protocol was applied for FACS, where 4×10^5^ A549 cells were plated per each well overnight in a 12-well plate. For FACS, instead of formaldehyde fixation, following incubation with the secondary antibody cells were detached using 10 mM EDTA, washed and centrifuged at 500 x g for 5 min, and subsequently analyzed by the Cytoflex flow cytometer. At least 5,000 events were recorded. FCS Express software was used for FACS data analysis.

### Ultra-high resolution scanning electron microscopy (UHR-SEM)

A549 cells were plated on indium tin oxide (ITO) coated coverslips (30-60 ohms). Coverslips were washed prior to cell plating as follows: 100% isopropanol, 70% ethanol, 100% ethanol, DDW, and moved to Dulbecco’s Modified Eagle Medium (DMEM). The coverslips were placed in 6-well plates and 1×10^6^ cells/well were seeded overnight. The plate with cells grown overnight was then chilled on ice and all the subsequent steps were performed at 4°C. Cells were incubated with NP (0.65μM) for 30min, then washed (3x cold PBS) and incubated with the relevant plasma sample diluted (1:50) in 1% BSA in PBS. Diluted plasma samples were ultra-centrifuged for 15 min, 120,000g at 4°C. The relevant coverslips were fixed using 2.5% PFA, 0.01% glutaraldehyde in 0.1M cacodylate buffer for 1h at RT. Coverslips were washed 3x with PBS, quenched 3x with 50 mM Tris, pH 8.0 and blocked using 5% BSA for 1h at RT. Coverslips (either fixed or non-fixed, as indicated) were then incubated with 18 nm colloidal gold-goat anti-human IgG (H+L) diluted (1:20) for 30 min. Cells were then rinsed and fixed using 2.5% glutaraldehyde in 0.2M cacodylate buffer, overnight at 4°C. The next day coverslips were washed 4x with ultra-pure water (UPW) and dehydrated gradually using 25%, 50%, 75% and 100% ethanol 2x washes with each ethanol concentration for 10 min at RT. After the ethanol washes, coverslips were left to air dry and coated with 1 nm iridium. Images were acquired using a ultra high-resolution scanning electron microscope (UHR-SEM, Zeiss Gemini 560), with Back Scattered electron Detector (BSD) at two different magnifications (10,000x and 50,000x). The imaging conditions were: accelerating voltage 5kV, aperture size 20 μm, focus 3.9mm and scanning speed 8.

### Cell-surface binding of FITC-labeled NP and its derivatives FACS

A549 cells were plated overnight in 12-well plates, 4×10^5^ cells/well. On the next day, the cells were chilled on ice and washed 3x with ice cold PBS, containing Ca^2+^ and Mg^2+^ All the subsequent steps were performed at 4°C. Labeled NP was serially diluted (0.13 μM, 0.65μM and 1.30 μM) and incubated with the cells for 1h, followed by 3x PBS wash. Cells were subsequently detached using 10 mM EDTA treatment and then washed twice with PBS (without Ca^2+^ and Mg^2+^) by centrifugation, 500 x g for 5 min. Cells were next analyzed by Cytoflex flow Cytometer. 10,000 events were recorded. Data analysis was conducted using the FCS Express software. A similar procedure was used for evaluation of binding and dissociation of NP derived constructs (the corresponding concentrations are as indicated in **Results**). For dissociation experiments, cells were thoroughly washed prior to measurement, while for binding experiments cells were immediately analyzed by flow cytometry after incubation with NP as described in **Results**.

### Enoxaparin treatment

For enoxaparin treatment, in each of the assays presented NP was added together with enoxaparin at the indicated concentrations and cells were processed as described in the corresponding procedure.

### RNA synthesis, labeling and EMSA

In vitro 242 nucleotide RNA (SARS-CoV-2, sequence ID OR193379.1, positions 19,763-20,004) was synthesized using primers 3345 (sense-forward primer, including the T7 RNA polymerase promoter, underlined: 5’-<u>CGAAATTAATACGACTCACTATAGGGACAG</u>GTTAATGTAGCATTTGAG) and 3383 (reverse primer 5’-ACTTGACCATCAACTC) and DNA template generated by TWIST Bioscience. The RNA was synthesized and purified as described in (67). To end label the RNA, 100 pmol of RNA were incubated with 10 units of calf intestinal phosphatase (CIP) in a 20 µl reaction at 37°C for 1h. The reaction was stopped by adding 20 µl of stop solution, heating at 70°C for 5 mins followed by loading onto a 6% acrylamide 8M urea gel. The RNA band was extracted from the gel followed by overnight incubation in RNA elution buffer (0.1 M sodium acetate, 0.1% SDS and 10 mM EDTA) at 4°C. The eluate was collected and purified by phenol : chloroform extraction, and precipitated with 1 ml of 100% ethanol and 20 µg of glycogen, followed by incubation at –80°C for 30 min or –20°C overnight. After centrifugation (13,000 rpm for 20 min at 4°C) the pellet was resuspended in DEPC. The dephosphorylated RNA (20 pmol) was then 5’-end labeled with T4-PNK at 37°C for 30 min. followed by phenol chloroform extraction. For EMSA, P_32_ labeled RNA was incubated with purified proteins, as indicated in **Figure 4** in buffer C (1x buffer C contains: 50 mM HEPES pH 7.5, 10 mM MgCl_2_, 100 mM NH_4_Cl, 15 mM DTT) at RT for 10 min, with a total reaction volume of 10 µl. Thereafter, 5 µl of native loading buffer containing 50% glycerol and 0.25% bromophenol blue was added and the mixture was analyzed in a 4% native gel, 0.5x TBE, at 150-volt, for 20 min in the cold room.

### Modeling of heparin binding to NP

Docking of heparin to CTD, NTD, and full NP (PDB ID 8FD5 (31)) was performed with the ClusPro server (https://cluspro.bu.edu) (27) using heparin as a ligand in the advanced settings. The resulting 8 top-scoring models were downloaded and visualized with ChimeraX v1.5 (68). The surface of the protein was colored either by conservation (calculated with the ConSurf server, https://consurf.tau.ac.il (69)) or by surface electrostatics, calculated by ChimeraX.

## Supplementary Data

Structural models of heparin docked to NTD, CTD and full-length NP, as well as a Chimera session used to generate **Figures 3B,C** are provided as a zipped file.

## Acknowledgements

We thank Prof. Hervé Bercovier and Prof. Mark Saper for helpful discussions and the critical reading of the manuscript. We would like to thank The Edmond and Benjamin de Rothschild Foundation for their generous support. This work was supported, in whole or in part, by the Israel Science Foundation, founded by the Israel Academy of Science and Humanities (grant numbers 338/19 to AR, and 3933/19 and 301/21 to O.S.-F.). J.K.V. is supported by a Marie Sklodowska-Curie European Training Network Grant #860517 (Ubimotif). We are thankful to the Hadassah blood bank members. We would like to especially thank all our cohort participants and to blood bank donors for their readiness and collaboration.

