## Supplementary Figures and Tables for "Binding of SARS-CoV-2 nucleocapsid protein to uninfected epithelial cells induces antibody-mediated complement deposition"

- Supplementary Figures 1-5
  - Figure S1 (accompanies Figure 2)
  - Figure S2 (accompanies Figure 3B,C)
  - Figure S3 (accompanies Figure 4F)
  - Figure S4 (accompanies Figure 5B,C)
  - Figure S5 (accompanies Figure 6E)
- Supplementary Tables 1-3
  - Table S1: Excel sheet
  - Table S2: Crystal structure details
  - Table S3: Primers used in this study
- Supplementary Datafiles (zip file)
  - Structural models of NTD-heparin, CTD-heparin, full NP-heparin complexes (Figure 3B,C and Supplementary Figure S3)
  - Chimera session used for generating Figure 3C

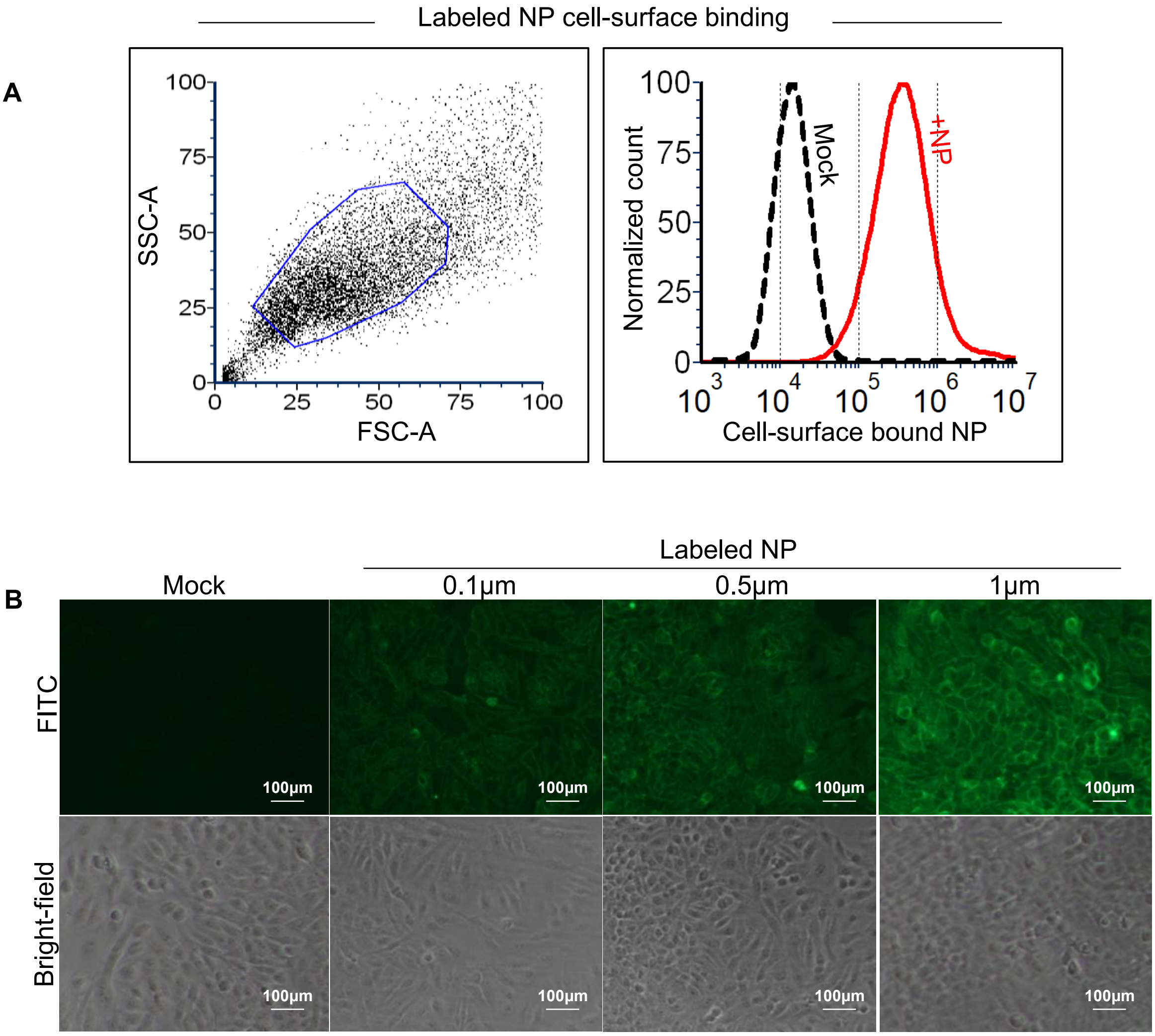


### Figure S1 (accompanies Figure 2)

**A**: The gating strategy: forward scatter area [FSC-A] *versus* side scatter area [SSC-A] was used to select cells (left). The histogram presents the cell-surface bound NP *versus* mock (right). **B:** Immunofluorescence microscopy of live HeLa cells showing concentration dependent binding of NP. Cells were incubated at 4°C with FITC-labeled NP at different concentrations (0, 0.1, 0.5, and 1 μM), and subsequently washed and formaldehyde fixed. Cells were imaged by fluorescence microscopy (20x LWD objective, Nikon, Eclipse T2i). Lower panels show the corresponding transmitted light microscopic image (phase contrast) to visualize cells. Scale bar represents 100 μm.

A

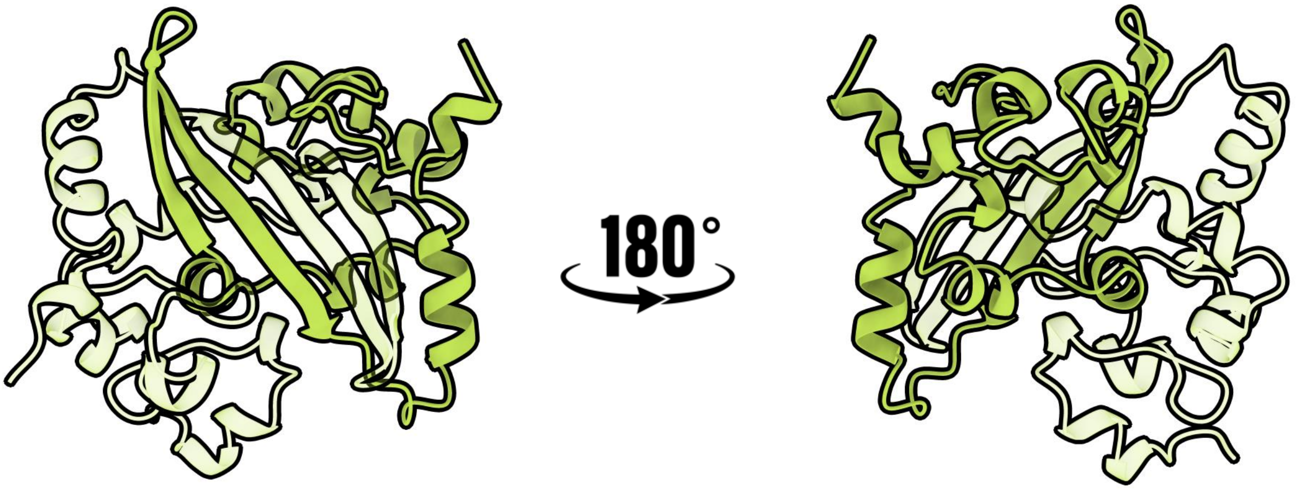


B

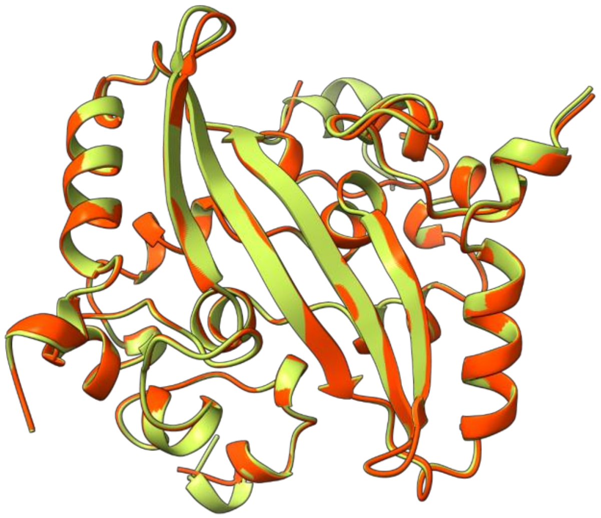

### Figure S2 (accompanies Figure 3B,C)

C

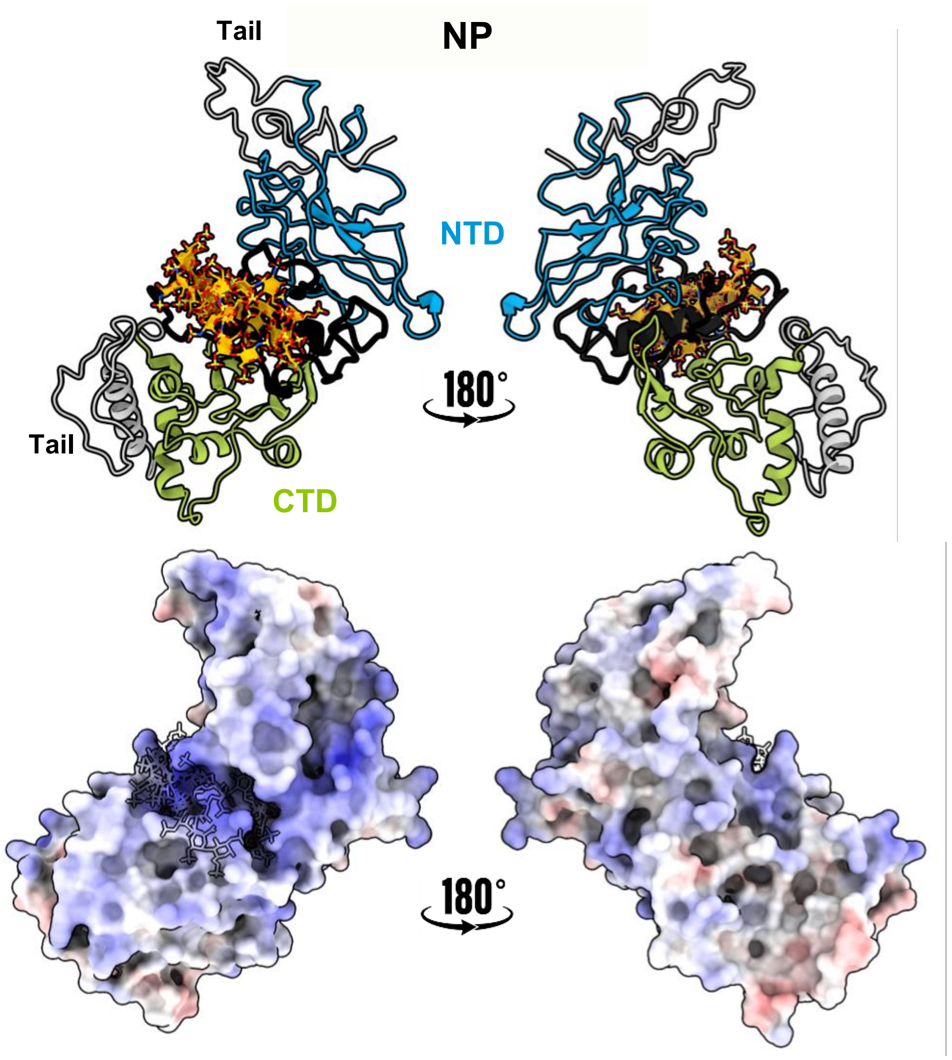


### Figure S2 (accompanies Figure 3B,C), continued

**A:** The CTD structure solved in this study (PDB ID 8R6E; see **Supplementary Table S2** for more details). The CTD dimer is shown in cartoon representation, in two different orientations. The two domains are colored in dark and light green, respectively. **B:** The structure is very similar to other solved CTD structures: Structural alignment to another representative solved CTD structure (PDB ID 7C22 (20,26,30), red), RMSD=0.17Å. **C:** Model of NP full-length structure (8FD5) bound to heparin through its linker (colored in black). **Upper panel**: Cartoon representation in two different orientations. **Lower panel**: The same orientations as the upper panel but colored according to the electrostatic potential, highlighting the positive patch at the linker binding site.

### Figure S3

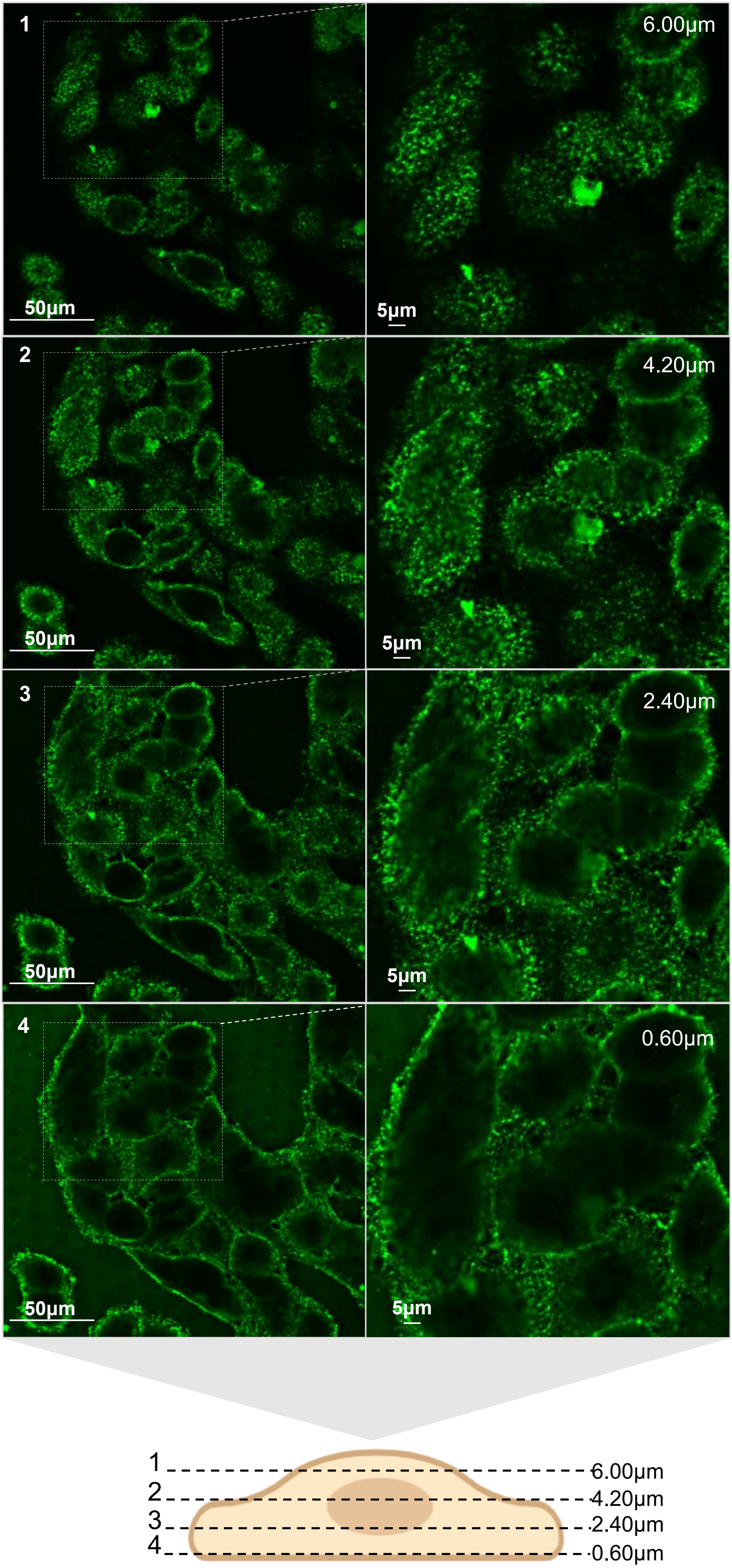
1
6.00μm
50μm
5μm
2
4.20μm
50μm
5μm
3
2.40μm
50μm
5μm
4
0.60μm
50μm
5μm
1
2
3
4
6.00μm
4.20μm
2.40μm
0.60μm

### Figure S3 (accompanies Figure 4F)

Confocal fluorescence microscopy of A549 cells that were incubated with FITC-labeled NP (0.5μM). Live cells were labeled at 4°C to prevent endocytosis and then fixed prior to confocal imaging (see **Methods**). Note the presence of cell surface patches, indicating clustered distribution of surface bound NP. Scale bars of 50μm and 5μm are shown**.** Schematic illustration at the bottom represents the shown focal planes of the presented field, the corresponding Z-focal distance is indicated at the top right corner. Images in the right panels represent digital zoom of the highlighted dotted region from the left panels.

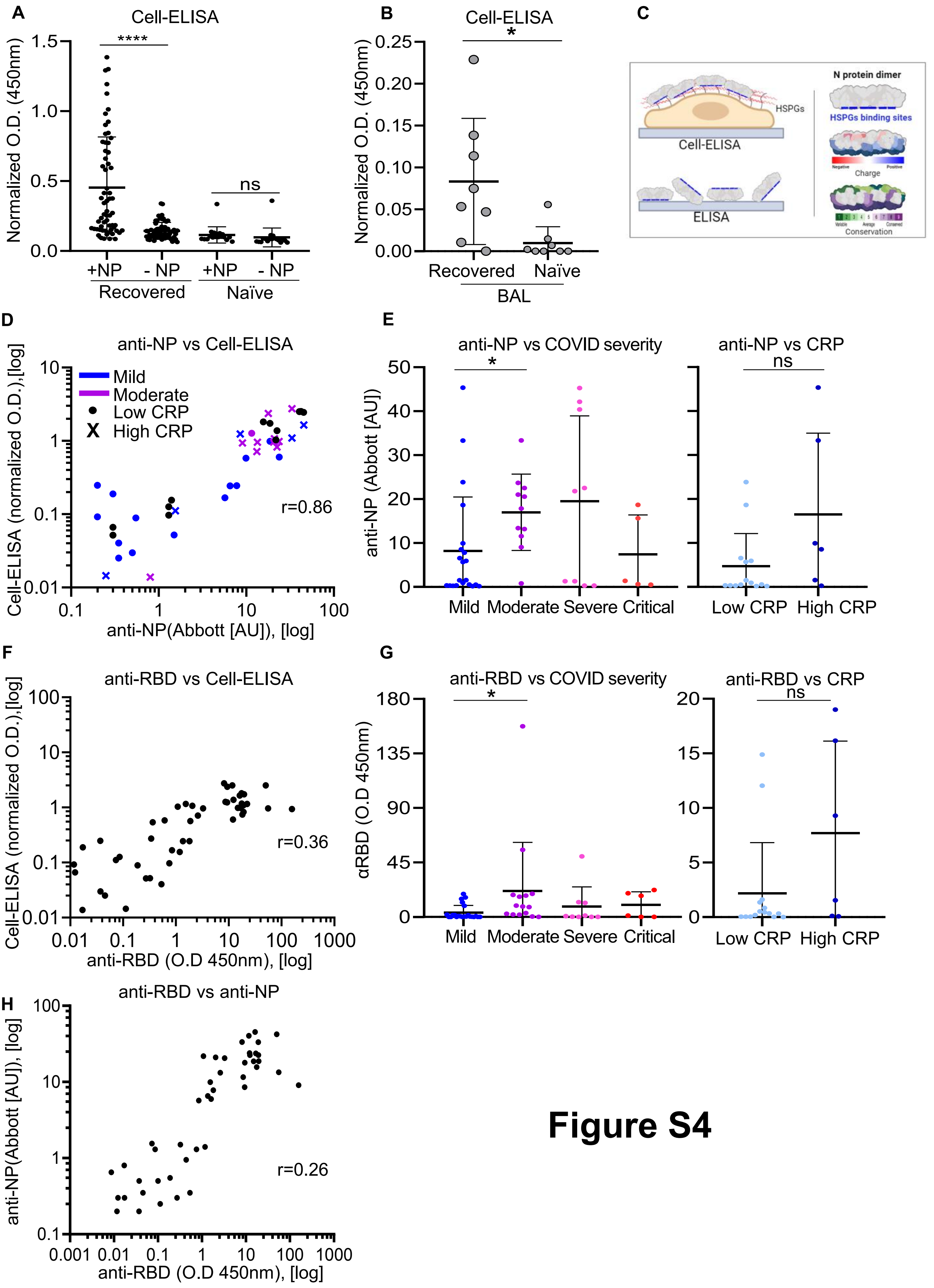
A
Cell-ELISA
Normalized O.D. (450nm)
****
ns
+NP
- NP
+NP
- NP
Recovered
Naïve
B
Cell-ELISA
*
Recovered
Naïve
BAL
C
N protein dimer
HSPGs
HSPGs binding sites
Cell-ELISA
Charge
ELISA
Conservation
D
anti-NP vs Cell-ELISA
Mild
Moderate
Low CRP
High CRP
r=0.86
Cell-ELISA (normalized O.D.),[log]
anti-NP(Abbott [AU]), [log]
E
anti-NP vs COVID severity
anti-NP vs CRP
ns
anti-NP (Abbott [AU])
Mild
Moderate
Severe
Critical
Low CRP
High CRP
F
anti-RBD vs Cell-ELISA
r=0.36
Cell-ELISA (normalized O.D.),[log]
anti-RBD (O.D 450nm), [log]
G
anti-RBD vs COVID severity
anti-RBD vs CRP
αRBD (O.D 450nm)
H
anti-RBD vs anti-NP
r=0.26
anti-NP(Abbott [AU]), [log]
anti-RBD (O.D 450nm), [log]

**Figure S4**

### Figure S4, continued

I

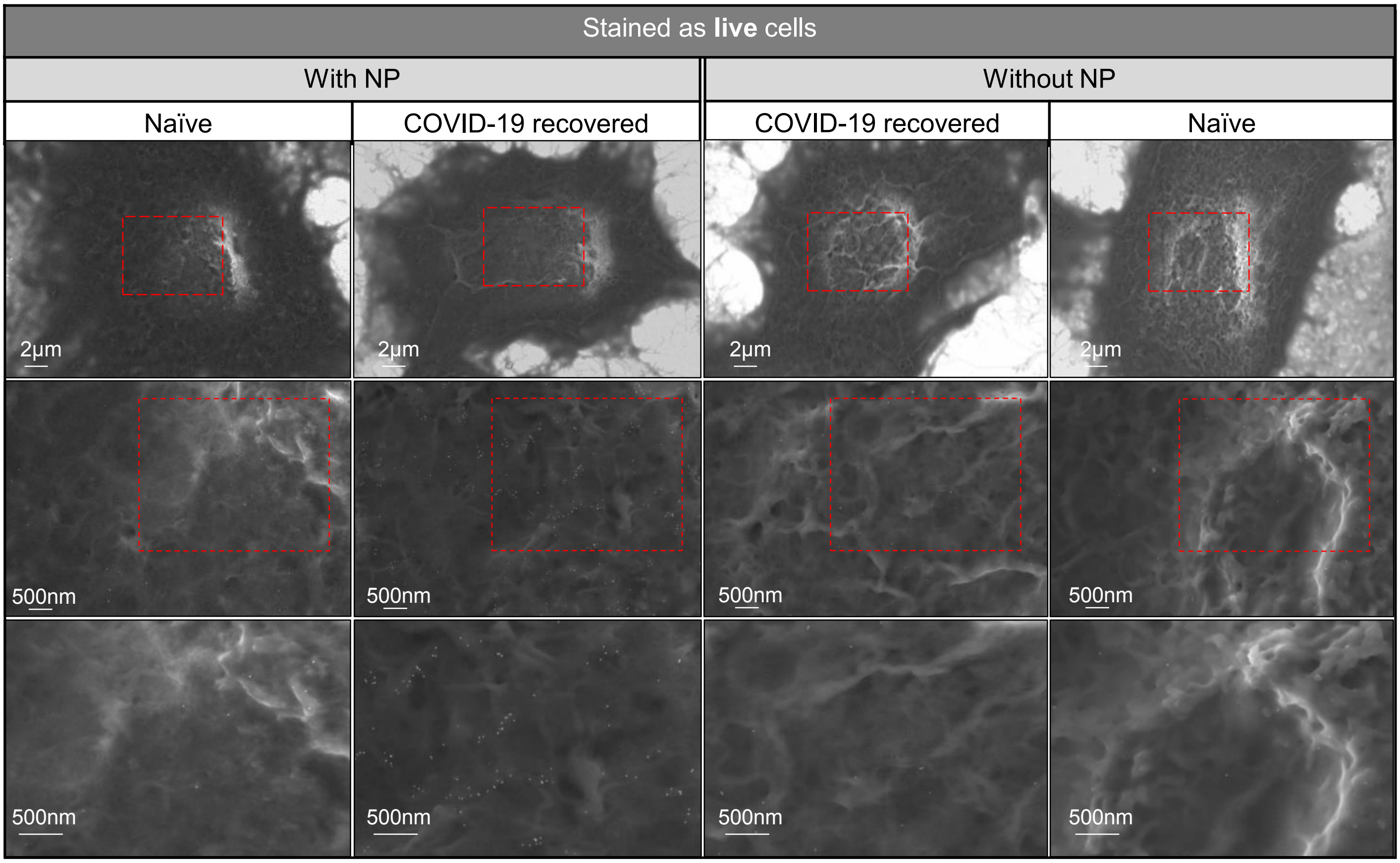

### Figure S4 (accompanies Figure 5B,C)

**A:** Control experiment to assess the specificity of anti-NP cell ELISA (see **Figure 5B**). Anti-NP cell ELISA was conducted on A549 cells using sera samples collected from COVID-19 recovered (n=54) or naïve (n=18) individuals. For evaluation of specificity, anti-NP cell ELISA was performed in either presence or in absence of NP (+NP and -NP, as indicated). The mean of three experimental replicas for each sample are plotted. The mean and standard deviation (SD) for each group of samples were calculated in GraphPad Prism. Student's two-tailed unpaired t-test was used to compare different groups: (*'****' P value* < 0.0001, recovered) and ('ns': *P* value= 0.382, naive) patient sera. **B:** Anti-NP cell ELISA using BAL fluid sample was performed using A549 cells coated with NP at 4°C that were subsequently incubated with BAL collected either from naïve (n=8) or from COVID-19 recovered individuals (n=8). The corresponding background control values (when NP was omitted from the assay) were subtracted from each presented measurement. The mean of three experimental replicas for each sample are plotted. The mean and standard deviation (SD) for recovered and naive groups were calculated in GraphPad Prism ('*': *P value* = 0.018). **C:** Schematic illustration of our model, highlighting the differences between anti-NP IgG cell ELISA versus classical anti-NP IgG assays (such as indirect ELISA) aimed to evaluate total anti-NP IgG. Note biased orientation of the antigen in the cell ELISA approach that only detects physiologically relevant epitopes of NP versus more random NP presentation in classical ELISA, see text for more details. **D:** Correlation between anti-NP IgG cell-ELISA values and total anti-NP IgG measured by ARCHITECT (see **Methods**) (n=46). Measurements are labeled according to COVID-19 severity classification, as indicated: blue for mild cases, purple for moderate cases, black are unclassified cases, 'X' marks high (>7.5) CRP values, while 'dot' marks low (<7.5) CRP values. CRP, C-Reactive Protein is a serum marker for inflammation (see **Methods**). **E:** Anti-NP results from ARCHITECT assay categorized by COVID-19 severity. Left panel (mild, moderate, severe and critical), n=46, '*': *P* value= 0.043. In the right panel the 'mild' category from the left panel was further sub-categorized based on CRP values (high>7.5, low<7.5), n=21, 'ns': *P value*=0.052. **F:** Pearson correlation analysis of anti-NP cell-ELISA *versus* total anti-RBD IgG measured by classical indirect ELISA (see **Methods**) (n=53), *r* = 0.36. **G:** Anti-RBD IgG measured by indirect ELISA categorized by COVID-19 severity (left, n=53. '*': *P value*= 0.042) and sub-categorized by CRP values (n=21, 'ns': *P valu*e= 0.067), as described in panel E. The corresponding background ELISA values were subtracted from each measurement (see panel **Supplementary Figure S4A** for details). **H:** Pearson correlation between anti-NP IgG results from ARCHITECT assay *versus* anti-RBD IgG, measured by indirect ELISA (see **Methods**), n=45, *r*=0.26. **I:** Immunogold Ultra-High Resolution Scanning Electron Microscopy (UHR-SEM) of A549 cells that were pre-incubated with full-length NP at 4°C (NP coating) and subsequently incubated with plasma samples collected from either naïve or COVID-19 recovered individuals, as indicated. Samples were then stained by goat-anti human IgG secondary antibodies conjugated to 18 nm colloidal gold particles for immunogold detection in SEM. All the staining steps were carried on live cells at 4°C to avoid endocytosis. To assess specificity of immunostaining, the coating step with NP was omitted (right panels without NP, as indicated). Scale bars represent: 2 µm and 500 nm, as indicated.

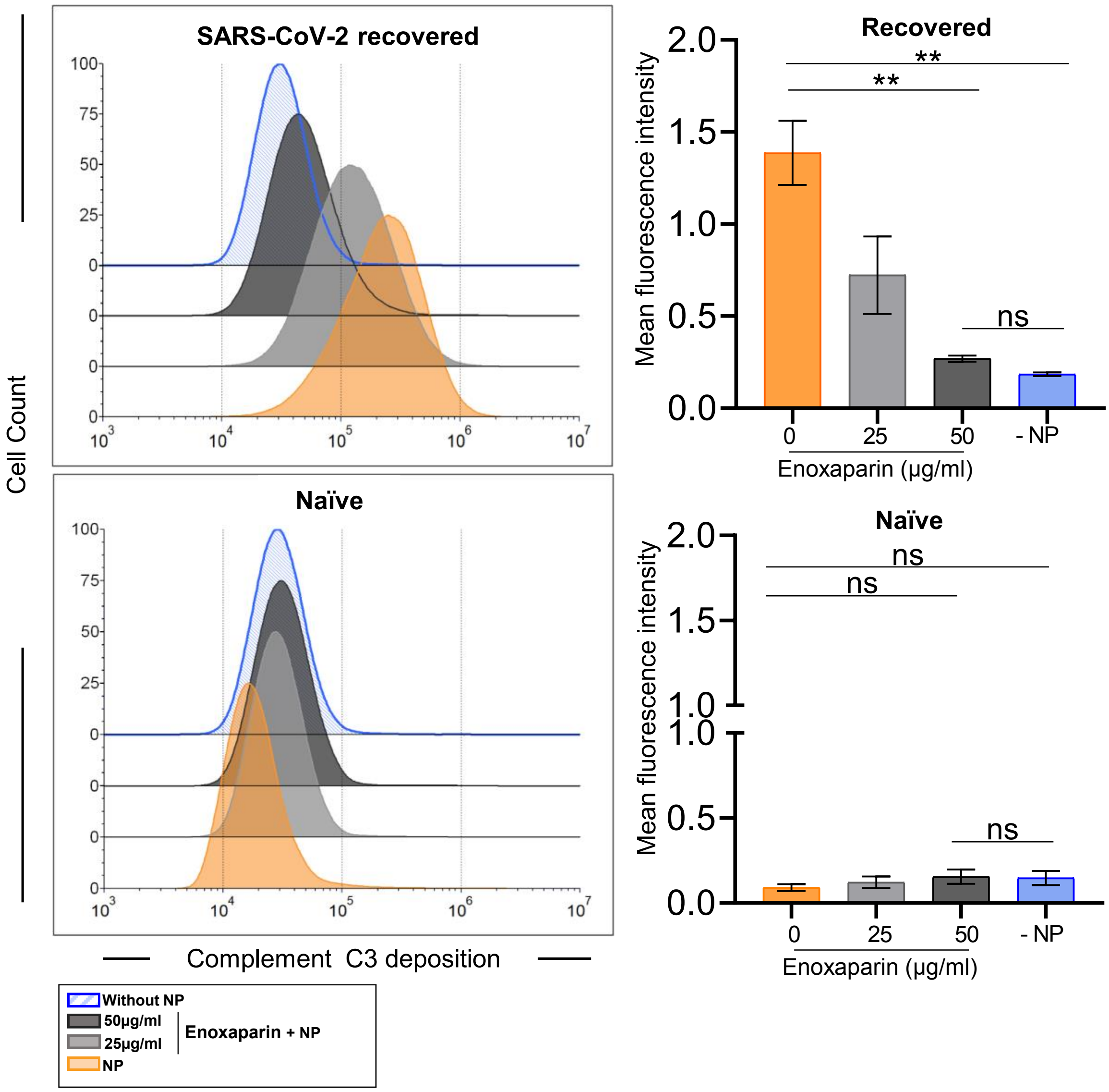


### Figure S5 (accompanies Figure 6E)

FACS, control experiments for the data presented in Figure 6E. A549 cells were either incubated with NP only, or co-incubated with NP and enoxaparin (25, 50 µg/ml). Subsequently, cells were incubated with plasma samples from either COVID-19 recovered or naïve individuals, followed by guinea pig complement. C3b deposition was evaluated using FITC conjugated goat anti-guinea pig complement, C3b. Samples where NP was omitted, were used to assess the specificity of immunostaining. The mean fluorescence intensity values of three independent replicates are plotted, showing mean +/- standard error (SEM). One-way analysis of variance (ANOVA) and Tukey's multiple comparisons test were used to compare between the different conditions. 'ns' indicates *P value* > 0.05, and '**' indicates *P value* < 0.01.

### Table S1: Cohort Excel file

**Details of the cohorts analyzed in this study (Excel file; accompanies Figures 1, 5, 6)**

Serum samples from the pre-vaccination era were collected from COVID-19 recovered and naïve individuals, (*) indicated samples collected in the pre-COVID era (n=231). Samples were categorized based on COVID-19 severity (mild, moderate, severe, and critical). Plasma samples were also collected from naïve or COVID-19 recovered individuals (n=8). An additional cohort of serum and Bronchoalveolar lavage (BAL) fluid samples was collected from COVID-19 recovered and naïve individuals (n=18), in the post-vaccination era. Individual values of C-reactive protein (CRP), white blood cells (WBC), ferritin, DDM, quantitative ELISA for anti-Np IgG (Abbott kit), anti-spike (liaison kit), ELISA anti-NTD (O.D.), ELISA anti-RBD (O.D.), cell-ELISA (normalized O.D.), CT and CXR to each sample are presented.

| | |
|---|---|
| Beamline | ESRF Beamline ID23-2 |
| Wavelength (Å) | 0.8731 |
| Space group | $P2_12_12_1$ |
| Unit Cell a, b, c (Å), α, β, γ (°) | 43.6, 47.4, 134.6, 90, 90, 90 |
| Resolution range (Å)[a] | 47.37 – 1.53 (1.55 - 1.53) |
| Total reflections [a] | 306,860 (13,074) |
| Unique reflections [a] | 43,189 (1,968) |
| Completeness (%)[a] | 99.6 (92.8) |
| Multiplicity [a] | 7.1 (6.6) |
| $R_{meas}$ (%) [a,b] | 16.6 (558.3) |
| $\langle I \rangle / \langle \sigma(I) \rangle$ [a] | 5.9 (0.4) |
| $CC_{1/2}$ [a,c] | 0.997 (0.291) |
| Wilson B-factor [d] (Å$^2$) | 29.6 |
| $R_{work}$ | 0.225 |
| $R_{free}$ | 0.267 |
| No. of protein monomers in a.u. | 2 |
| Number of atoms | |
| Macromolecules | 1923 |
| Solvent | 208 |
| Number of protein residues | 234 |
| RMS bond lengths (Å) | 0.008 |
| RMS bond angles (°) | 1.70 |
| Ramachandran favored (%)[e] | 98.7 |
| Ramachandran allowed (%) | 1.3 |
| Ramachandran outliers (%)[f] | 0.0 |
| Clashscore [e] | 4.3 |
| Average B-factor protein (Å$^2$) | 37.8 |
| Average B-factor solvent (Å$^2$) | 48.4 |
| RCBS PDB code | 8R6E |

[a]Values for the highest resolution shell are given in parentheses

[b] $R_{meas} = \Sigma_h[m/(m - 1)]^{1/2} \Sigma_i|I_{h,i} - \langle I_h \rangle| / \Sigma_h \Sigma_i I_{h,i}$

[c] $CC_{1/2}$ is defined in (Karplus & Diederichs, 2012).

[d] Wilson B-factor was estimated by SFCHECK. (Vaguine et al., 1999).

[e] The Ramachandran statistics and clashscore statistics were calculated using MOLPROBITY. (Williams et al., 2018).


Karplus and Diederichs (2012) Linking crystallographic model and data quality. *Science* **336**:1030-1033.

Tickle, Flensburg, Keller, Paciorek, Sharff, Vonrhein, and Bricogne (2018). STARANISO (Global Phasing Ltd.).

Vaguine, Richelle, and Wodak (1999). SFCHECK: a unified set of procedures for evaluating the quality of macromolecular structure-factor data and their agreement with the atomic model. *Acta Crystallogr. D Biol. Crystallogr.* **55**:191–205.

Williams, Headd, Moriarty, Prisant, Videau, Deis, Verma, Keedy, Hintze, Chen, Jain, Lewis, Arendall, Snoeyink, Adams, Lovell, Richardson, Richardson(2018). MolProbity: More and better reference data for improved all-atom structure validation. *Protein Sci*. **27**:293-315.


### Table S2

### Table S3: Primers used in this study

| Primer | Sequence (5'-3') | Comments |
|---|---|---|
| 458 (F)<br>459 (R) | TATGGAGAATCTTTACTTTCAGGGGATGTCTGATAATGGACCCCAAAATC<br>GGCTTTGTTAGCAGCCGGATCCTCGAGTTAGGCCTGAGTTGAGTCAGC | Amplify NP full length for pET15B |
| 462 (F)<br>463 (R) | TATGGAGAATCTTTACTTTCAGGGGCGGCCCCAAGGTTTACCCAATAATAC<br>GGCTTTGTTAGCAGCCGGATCCTCGAGtcaTTCTGCGTAGAAGCCTTTTGG | Amplify NP 41-174 (NTD) for pET15B |
| 464 (F)<br>465 (R) | TATGGAGAATCTTTACTTTCAGGGGACTAAGAAATCTGCTGCTGAG<br>GGCTTTGTTAGCAGCCGGATCCTCGAGtcaTGGGAATGTTTTGTATGCGTC | Amplify NP 247-364 (CTD) for pET15B |
| 462 (F)<br>465 (R) | TATGGAGAATCTTTACTTTCAGGGGCGGCCCCAAGGTTTACCCAATAATAC<br>GGCTTTGTTAGCAGCCGGATCCTCGAGtcaTGGGAATGTTTTGTATGCGTC | Amplify NP 41-364 (NTD-IDL-CTD) for pET15B |
| (F)<br>465 (R) | GAAggtagcggtagcCTGGTGCCGCGCGGCAGCgggtcagggtcaaccAAGAAGAGCGCGGCAG<br>GGCTTTGTTAGCAGCCGGATCCTCGAGtcaTGGGAATGTTTTGTATGCGTC | Amplify GSGSLVPRGSGSGS-**CTD** |
| 462 (F)<br>(R) | TATGGAGAATCTTTACTTTCAGGGGCGGCCCCAAGGTTTACCCAATAATAC<br>ggttgaccctgacccGCTGCCGCGCGGCACCAGgctaccgctaccTTCAGCATAAAATCCTTTG | Amplify **NTD**-GSGSLVPRGSGSGS |
